# Comprehensive analysis of T cell immunodominance and immunoprevalence of SARS-CoV-2 epitopes in COVID-19 cases

**DOI:** 10.1101/2020.12.08.416750

**Authors:** Alison Tarke, John Sidney, Conner K Kidd, Jennifer M. Dan, Sydney I. Ramirez, Esther Dawen Yu, Jose Mateus, Ricardo da Silva Antunes, Erin Moore, Paul Rubiro, Nils Methot, Elizabeth Phillips, Simon Mallal, April Frazier, Stephen A. Rawlings, Jason A. Greenbaum, Bjoern Peters, Davey M. Smith, Shane Crotty, Daniela Weiskopf, Alba Grifoni, Alessandro Sette

**Affiliations:** Center for Infectious Disease and Vaccine Research, La Jolla Institute for Immunology (LJI), La Jolla, CA 92037, USA; Department of Internal Medicine and Center of Excellence for Biomedical Research (CEBR), University of Genoa, Genoa, 16132, Italy; Department of Medicine, Division of Infectious Diseases and Global Public Health, University of California, San Diego (UCSD), La Jolla, CA 92037, USA; Institute for Immunology and Infectious Diseases, Murdoch University, Perth, Western Australia, Australia

## Abstract

T cells are involved in control of SARS-CoV-2 infection. To establish the patterns of immunodominance of different SARS-CoV-2 antigens, and precisely measure virus-specific CD4^+^ and CD8^+^ T cells, we studied epitope-specific T cell responses of approximately 100 convalescent COVID-19 cases. The SARS-CoV-2 proteome was probed using 1,925 peptides spanning the entire genome, ensuring an unbiased coverage of HLA alleles for class II responses. For HLA class I, we studied an additional 5,600 predicted binding epitopes for 28 prominent HLA class I alleles, accounting for wide global coverage. We identified several hundred HLA-restricted SARS-CoV-2-derived epitopes. Distinct patterns of immunodominance were observed, which differed for CD4^+^ T cells, CD8^+^ T cells, and antibodies. The class I and class II epitopes were combined into new epitope megapools to facilitate identification and quantification of SARS-CoV-2-specific CD4^+^ and CD8^+^ T cells.

## INTRODUCTION

As of November 2020, SARS-CoV-2 infections are associated with 1.2 million deaths and over 48 million cases worldwide, and over 9 million cases in the United States alone (https://coronavirus.jhu.edu/map.html). The severity of the associated Coronavirus Disease 2019 (COVID-19) ranges from asymptomatic or mild self-limiting disease, to severe pneumonia and acute respiratory distress syndrome (WHO; https://www.who.int/publications/i/item/clinical-management-of-covid-19). We and others have started to delineate the role of SARS-CoV-2-specific T cell immunity in COVID-19 clinical outcomes (Altmann and Boyton, 2020; Braun et al., 2020; Grifoni et al., 2020; Le Bert et al., 2020; Meckiff et al., 2020; Rydyznski Moderbacher et al., 2020; Sekine et al., 2020; Weiskopf et al., 2020). A growing body of evidence points to a key role for SARS-CoV-2-specific T cell responses in COVID-19 disease resolution and modulation of disease severity (Rydyznski Moderbacher et al., 2020; Schub et al., 2020; Weiskopf et al., 2020). Milder cases of acute COVID-19 were associated with coordinated antibody, CD4^+^ and CD8^+^ T cell responses, whereas severe cases correlated with a lack of coordination of cellular and antibody responses, and delayed kinetics of adaptive responses (Rydyznski Moderbacher et al., 2020; Weiskopf et al., 2020).

To date, most studies have utilized pools of predicted or overlapping peptides spanning the sequence of different SARS-CoV-2 antigens (Altmann and Boyton, 2020; Grifoni et al., 2020; Meckiff et al., 2020; Peng et al., 2020; Rydyznski Moderbacher et al., 2020; Schub et al., 2020; Sekine et al., 2020; Weiskopf et al., 2020), but the exact T cell epitopes and immunodominant antigen regions have not been comprehensively determined. Several studies have mapped different epitopes or the corresponding T cell receptors (TCRs) (Ferretti et al., 2020; Keller et al., 2020; Le Bert et al., 2020; Nelde et al., 2020; Peng et al., 2020; Snyder et al., 2020), but have been biased in their approach, due to sampling only a limited number of cells (Ferretti et al., 2020; Keller et al., 2020; Sekine et al., 2020), using HLA predictions focused on a limited number of allelic variants not representative of the majority of the human population (Ferretti et al., 2020; Keller et al., 2020), utilizing *in vitro* re-stimulation protocols (Keller et al., 2020; Nelde et al., 2020), or detecting responses mediated by only a few cytokines, potentially largely underestimating total responses (Keller et al., 2020; Le Bert et al., 2020).

Defining a comprehensive set of epitope specificities is important for several reasons. First, it allows us to determine whether within different SARS-CoV-2 antigens certain regions are immunodominant. This will be important for vaccine design, so as to ensure that vaccine constructs include not only regions targeted by neutralizing antibodies, such as the receptor binding domain (RBD) in the spike (S) region, but also include regions capable of delivering sufficient T cell help, and are suitable targets of CD4^+^ T cell activity. Second, a comprehensive set of epitopes helps define the breadth of responses, in terms of the average number of different CD4^+^ and CD8^+^ T cell SARS-CoV-2 epitopes generally recognized by each individual. This is key because some reports have described a T cell repertoire focused on few viral epitopes (Ferretti et al., 2020), which would be concerning for potential viral escape from immune recognition via accumulated mutations that can occur during replication or through viral reassortment. Third, a comprehensive survey of epitopes restricted by a set of different HLAs representative of the diversity present in the general population is important to ensure that results obtained are generally applicable across different ethnicities and racial groups, and also to lay the foundations to examine the potential associations of certain HLAs with COVID-19 severity. Finally, the definition of the epitopes recognized in SARS-CoV-2 infection is relevant in the context of the debate on the potential influence of SARS-CoV-2 cross-reactivity with endemic “Common Cold” Coronaviruses (CCC) (Braun et al., 2020; Le Bert et al., 2020). Several studies have defined the repertoire of SARS-CoV-2 epitopes recognized in unexposed individuals (Braun et al., 2020; Mateus et al., 2020; Nelde et al., 2020), but the correspondence between that repertoire and the epitope repertoire elicited by SARS-CoV-2 infection has not been evaluated.

In this study, we report a comprehensive map of epitopes recognized by CD4^+^ and CD8^+^ T cell responses across the entire SARS-CoV-2 viral proteome. Importantly, these epitopes have been characterized in the context of a broad set of HLA alleles using a direct *ex vivo*, cytokine-independent, approach.

## RESULTS

### Characteristics of the study participants

To broadly define the pattern of immunodominance and epitope recognition associated with SARS-CoV-2 infection, we studied PBMC samples from 99 adult convalescent COVID-19 donors. Their age ranged from 19 to 91 years (median 41), with a gender ratio of about 2M:3F (Male 41%; Female 59%). Ethnic breakdown was reflective of the demographics of the local enrolled population. Samples were obtained 3 to 184 days post-symptom onset (median 67 days). Peak COVID-19 disease severity was representative of the distribution observed in the general population to date (mild 91%, moderate 2%, severe and critical 7%) (**Table 1**).

**Table 1.** Characteristics of donor cohort utilized in the protein screen.

|  | **COVID-19 (n = 99)** |
| --- | --- |
| **Age (years)** | 19-91 [median = 41, IQR = 23] |
| **Gender** |  |
| **Male (%)** | 41% (42/99) |
| **Female (%)** | 59% (58/99) |
| **Sample Collection Date** | Mar-Sep 2020 |
| **SARS-CoV-2 PCR** | Positive = 100% (78/78)<br>Not tested= 21% (21/99) |
| **S RBD IgG** | 98% (97/99) |
| **Peak disease Severity<sup>a</sup>** |  |
| **Mild** | 91% (90/99) |
| **Moderate** | 2% (2/99) |
| **Severe** | 1% (1/99) |
| **Critical** | 6% (6/99) |
| **Race-Ethnicity** |  |
| **White- not Hispanic or Latino** | 76% (76/99) |
| **Hispanic or Latino** | 12% (12/99) |
| **Asian** | 5% (5/99) |
| **American Indian/Alaska Native** | 1% (1/99) |
| **Native Hawaiian or other Pacific Islander** | 0% (0/99) |
| **Black or African American** | 2%(2/99) |
| **More than one race** | 3% (3/99) |
| **Not reported** | 0% (0/99) |
| **Days Post Symptom Onset at Collection<sup>b</sup>** | 3-184 (108/108) [Median = 67, IQR = 48] |

<sup>a</sup>According to WHO criteria.
<sup>b</sup>Multiple visits for the same donor have been analyzed.

SARS-CoV-2 infection was determined by PCR-based testing during the acute phase of infection, if available (79% of the cases), and/or verified by plasma SARS-CoV-2 S protein RBD IgG ELISA (Stadlbauer et al., 2020) using plasma from convalescent phase blood draws. All donors were seropositive at the time of blood donation, with the exception of two mildly symptomatic donors with positive PCR results from the acute phase of illness, but seronegative results at time of blood donation (at 55 and 148 days post-symptom onset (PSO), respectively).

All donors were HLA typed at both class I and class II loci (**Table S1**). The HLA class I and II alleles frequently observed in the enrolled cohort were largely reflective of what is found in the worldwide population, as reported by the Allele Frequency Net Database (Gonzalez-Galarza et al., 2020), and as retrieved from the Immune Epitope Database’s (IEDB; www.iedb.org) population coverage tool (Bui et al., 2006; Dhanda et al., 2019) (**Fig. S1**). Of the 20 different HLA class I alleles with phenotypic frequencies >5% in our cohort, 15 (75%) are also present in the most common and representative class I alleles in the worldwide population (Paul et al., 2013) (**Fig. S1A-B**). Likewise, of the 34 different HLA class II alleles with phenotypic frequencies >5% in our cohort, 26 (76%) are also present in the worldwide population with frequencies >5%. These alleles correspond to 16 of the 27 (59%) alleles included in a reference panel of the most common and representative class II alleles in the general population (Greenbaum et al., 2011)(**Fig. S1D-F**). In conclusion, our cohort is largely representative of the HLA allelic variants commonly expressed worldwide.

### Pattern of antigen immunodominance in CD4+ and CD8+ T cell responses to SARS-CoV-2 antigens

To study adaptive immune responses in COVID-19 convalescent donors, we previously utilized T cell receptor (TCR) dependent Activation Induced Marker (AIM) assays to quantify SARS-CoV-2—specific CD4^+^ and CD8^+^ T cells utilizing the combination of markers OX40^+^CD137^+^ and CD69^+^CD137^+^ for CD4^+^ and CD8^+^ T cell, respectively (Grifoni et al., 2020; Mateus et al., 2020; Weiskopf et al., 2020). To define the global pattern of immunodominance in the study cohort, we tested PBMC from each donor with sets of overlapping peptides spanning the various SARS-CoV-2 proteins, as previously described (Grifoni et al., 2020b) (**Fig. 1A-B).**These data also defined the specific viral antigens recognized by each donor, and therefore highlight the specific antigens/donor pairs suitable for further epitope identification studies, as shown in **Fig. 1C and E.**

**Fig. 1.**
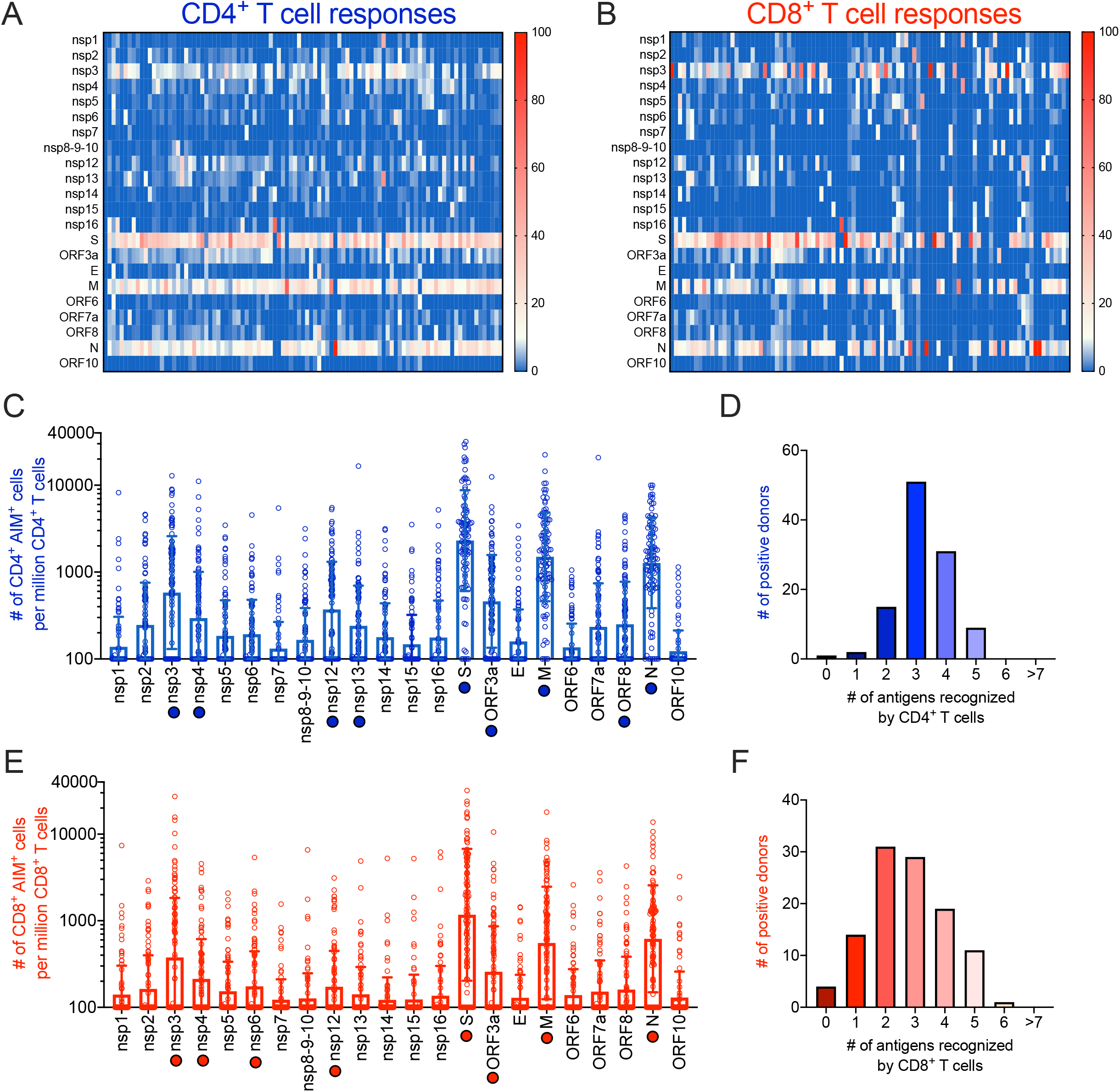
SARS-CoV-2-specific T cell reactivity per protein. Heatmaps of T cell reactivity across the entire SARS-CoV-2 proteome and as a function of the donor tested are shown for CD4^+^ (**A**) and CD8^+^ (**B**) T cells. Immunodominance at the ORF/antigen level and breath of T cell responses are shown for CD4^+^ (**C**) and CD8^+^ (**E**) T cells. Data are shown as geometric mean ± geometric SD. The numbers of donors recognizing one or more antigens with a response >10%, normalized per donor to account for the differences in magnitude based on days PSO, are shown for CD4^+^ (**D**) and CD8^+^ (**F**) T cells. Empty blue and red circles represent CD4^+^ and CD8^+^ T cell reactivity per protein, respectively. Filled blue and red circles highlight the immunodominant antigens recognized by CD4^+^ and CD8^+^ T cells, respectively.

For each SARS-CoV-2 protein antigen (**Table 2**) we recorded the % of donors in which a positive response was detected and the total response counts (positive cells/million detected in the AIM assay). This information was used to tabulate the percentage of the total response ascribed to each protein, and calculate the cumulative coverage provided by the most immunodominant proteins.

**Table 2.**
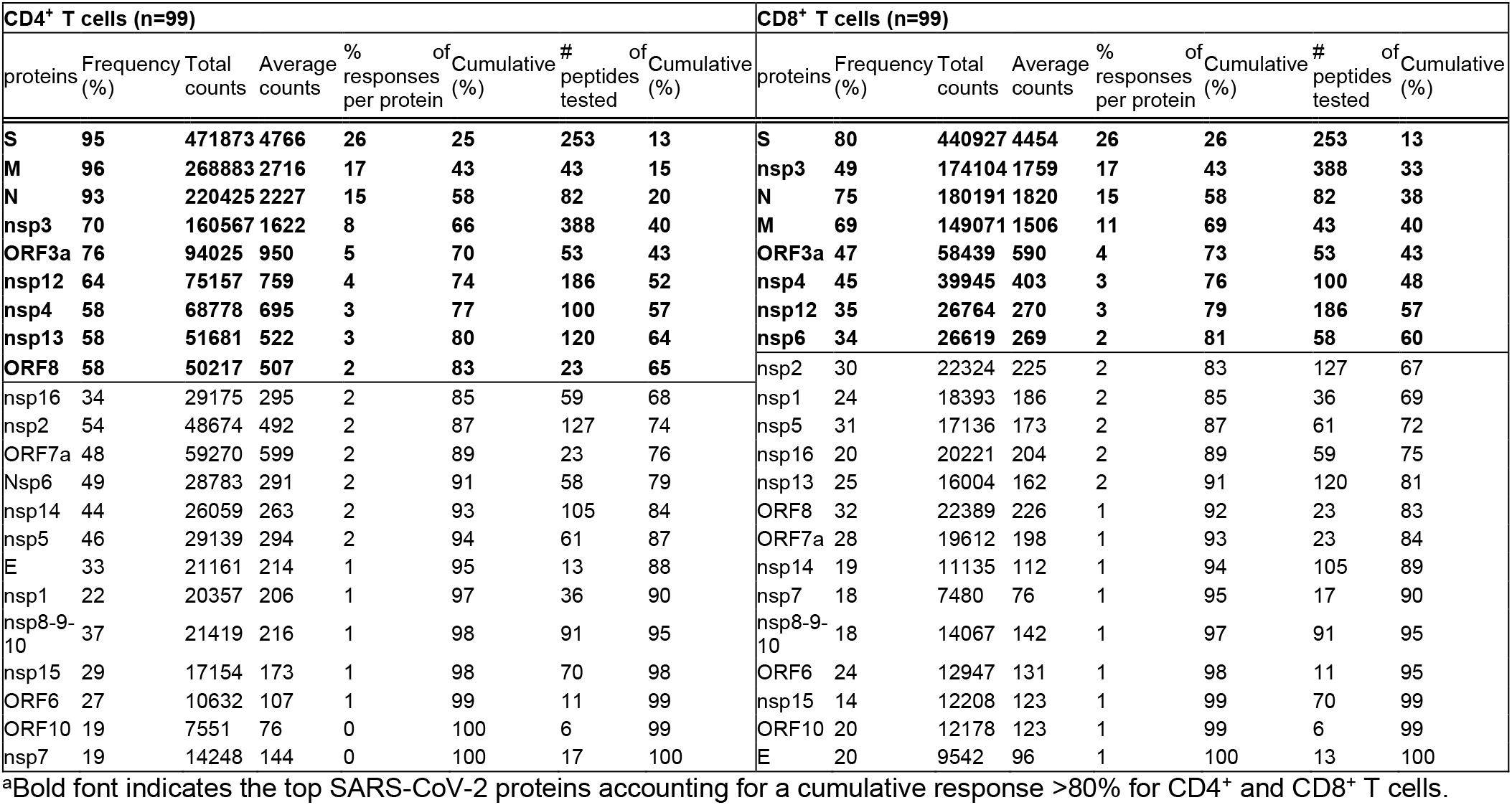
Summary of SARS-CoV-2-specific T cell reactivity as a function of the most immunodominant proteins^a^

For CD4^+^ T cell responses, 9 viral proteins (non-structural protein (nsp) 3, nsp4, nsp12, nsp13, S, ORF3a, Membrane (M), ORF8, and Nucleocapsid (N)) accounted for 83% of the total response. In the context of CD8^+^ T cell responses, 8 viral proteins (nsp3, nsp4, nsp6, nsp12, S, ORF3a, M, and N) accounted for 81% of the total response. These results confirmed the pattern previously observed with a more limited (n=20) number of COVID-19 patients (Grifoni et al., 2020b) and highlight a broad pattern of immunodominance, where 8-9 antigens are required to cover 80% of the response.

We further evaluated the number of antigens recognized in each of the individual donors analyzed. To this end, we focused on antigens associated with a sizeable response, arbitrarily defined herein as those antigens individually accounting for at least 10% of the total response. We found that per donor an average of 3.2 and 2.7 proteins were recognized by 10% or more of the total CD4^+^ and CD8^+^ SARS-CoV-2-specific T cells, respectively (**Fig. 1D and F**).

### Functional consequences of SARS-CoV-2—specific CD4+ T cell responses directed against different antigens

We next investigated whether the recognition of different SARS-CoV-2 antigens by CD4^+^ T cells correlated with functional antibody and/or CD8^+^ T cell responses. Consistent with the wide range of blood collection time points (day PSO) and peak disease severity in the COVID-19 donor cohort, we observed a wide range of RBD IgG responses (**Fig. 2A**). Positive correlations were observed between the RBD IgG antibody titers and the CD4^+^ T cell responses directed against the S (R= 0.2223, p= 0.0270), M (R= 0.2117, p= 0.0354), and ORF8 (R= 0.1982, p= 0.0492) antigens; no correlation was observed for the other viral antigens **(Fig. 2B-E)**.

**Fig. 2.**
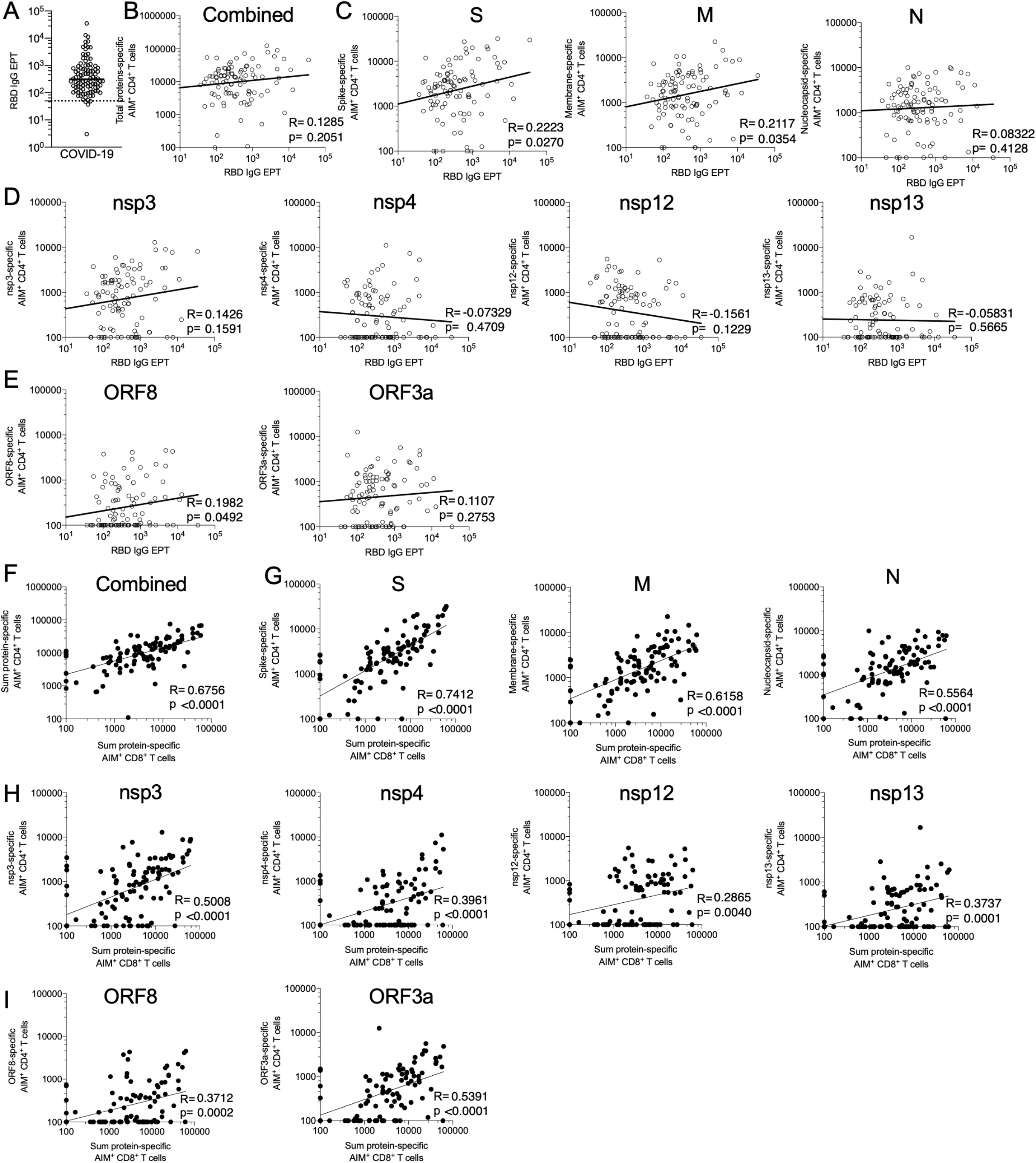
SARS-CoV-2-specific CD4^+^ T cell reactivities and their correlation with antibody production and CD8^+^ T cell reactivity. RBD IgG serology is shown for all the donors of this cohort (**A**). Serology data of panel **A**are correlated with CD4^+^ T cell reactivities specific against all combined proteins (**B**), structural proteins S, M, and N (**C**), non-structural proteins nsp3, nsp4, nsp12 and nsp13 (**D**), and ORF8 and ORF3a (**E**). The total CD8^+^ T cell reactivity is correlated with the total CD4^+^ T cell reactivity (**F**) and the CD4^+^ T cell reactivity against structural proteins S, M, and N (**G**), non-structural proteins nsp3, nsp4, nsp12 and nsp13 (**H**), and ORF8 and ORF3a (**I**). Empty and filled circles represent correlation between CD4^+^ T cell reactivity and serology or CD8^+^ T cell reactivity, respectively. All analyses were performed using Spearman correlation.

We then further examined the correlation between total CD8^+^ T cell responses and total CD4^+^ T cell responses and a strong correlation was observed (R= 0.6756, p <0.0001) (**Fig. 2F**). This correlation was retained when considering CD4^+^ T cell responses against the 9 most immunodominant antigens individually (**Fig. 2G-I)**. These data overall suggest that the CD4^+^ T cell response against all dominant antigens is potentially relevant in terms of providing helper function for CD8^+^ T cell-specific responses. This might reflect that T cell responses correlate with gene expression. S, N, and M may be immunodominant because of the very high gene expression for each (Xie et al., 2020). In this context, it is perhaps surprising that a strong CD4^+^ and CD8^+^ T cell response was elicited by nsp3, which is not known to be expressed at high levels (Xie et al., 2020).

### SARS-CoV-2 peptides and epitope screening strategy

The analysis of the SARS-CoV-2 proteome summarized above identified the major viral antigens accounting for 80% or more of the total CD4^+^ and CD8^+^ T cell response. These antigens were then introduced into the epitope screening pipeline (**Fig. S2**). Since class II epitope prediction is not as robust as class I prediction (Peters et al., 2020), and because of the high degree of overlap in binding capacity of different HLA class II alleles, to determine CD4^+^ T cell reactivity in more detail we followed a comprehensive and unbiased approach based on the use of complete sets of overlapping peptides spanning each antigen, and composition of antigen-specific peptide pools. Positivity was defined as net AIM^+^ counts (background subtracted by the average of triplicate negative controls) >100 and a Stimulation Index (SI) >2, as previously described (da Silva Antunes et al., 2020). Positive peptide pools were deconvoluted to identify the specific 15-mer peptide(s) recognized. For large proteins, such as S, an intermediate “mesopool” step was used to optimize use of reagents.

In parallel, we synthesized panels of predicted HLA class I binders for the 28 most common allelic variants (**Table S2**), as described in the methods section. The top two hundred predicted peptides were synthesized for each allele, leading to 5,600 predicted HLA binders in total. To identify CD8^+^ T cell epitopes, we tested individual peptides derived from the specific antigen(s) recognized by CD8^+^ T cells of individual donors and that were predicted to bind the HLA class I alleles expressed by the respective donor. (**Fig. 1B and E**). To quantify the population coverage provided by the HLA class I alleles selected for study, we tabulated the fraction of the donor cohort studied where allele matches were identified for 0, 1, 2, 3 or 4 of the respective HLA A and B alleles expressed by the donor. We found that 98% of the participants in our cohort were covered by at least one allele, 91% by 2 or more; 74% were covered by 3 or more of the alleles in our panel (**Fig. S1C**). As shown in **Table 2**, focusing on the 8 most dominant SARS-CoV-2 antigens for the purpose of epitope identification allowed mapping of 80% or more of the response, while screening only 35-40% of the total peptides.

To broadly identify T cell epitopes recognized in a cytokine-independent manner, we used the AIM assay mentioned above (Grifoni et al., 2020; Reiss et al., 2017). Examples of gating strategies, pool deconvolution and epitope identification for both CD4^+^ and CD8^+^ T cell responses are shown in **Fig. S2B.**AIM^+^ cell counts were calculated per million CD4^+^ or CD8^+^ T cells, respectively.

### Summary of CD4^+^ T cell epitope identification results

To identify specific CD4^+^ T cell epitopes, we deconvoluted peptide pools corresponding to antigens previously identified as positive for CD4^+^ T cell activity in each specific donor (**Fig. 1A**). In instances where not all positive pools could be deconvoluted due to limited cell availability, peptide pools were selected for screening to ensure that each of the 9 major antigens was tested in at least 10 donors. Overall, we were able to test each peptide for these antigens in a median of 13 donors (range 10 to 17). Each donor was previously determined to be positive for CD4^+^ T cell responses to that specific antigen.

Taken together, a total of 280 SARS-CoV-2 CD4^+^ T cell epitopes were identified, including 3 nsp16 (this protein was not included in the top proteins studied) epitopes identified in parallel experiments in 2 donors (**Table S3**). We found that each donor responded to an average of 3.2 viral antigens (**Fig. 1D**), and 5.9 CD4^+^ T cell epitopes were recognized per antigen for the top 80% most immunodominant antigens (data not shown). For each epitope/responding donor combination, potential HLA restrictions were also inferred based on the predicted HLA binding capacity of the epitope for the HLA alleles present in the respective responding donor (listed in **Table S1**), as previously described (Mateus et al., 2020; Voic et al., 2020).

### HLA binding capacity of dominant epitopes

A total of 109 of the 280 epitopes were recognized by 2 or more donors, accounting for 71% of the total response. The 49 most dominant epitopes, recognized in 3 or more donors, accounted for 45% of the total response (**Fig. S3A**).

Since dominant epitopes are associated with promiscuous HLA class II binding (Lindestam Arlehamn et al., 2013; Oseroff et al., 2010), defined as the capacity to bind multiple HLA allelic variants, we investigated the role of HLA binding in determining immunodominant SARS-CoV-2 epitopes. Specifically, we measured the *in vitro* binding capacity of the 49 most dominant epitopes (positive in 3 or more donors, as mentioned above) for a panel of 15 of the most common DR alleles using individual peptides and purified HLA class II molecules (Sidney et al., 2013). The results are provided in **Table S4**. It was noted that, in general, a good correlation was observed between predicted and measured binding (R= 0.6604, p<0.0001; **Fig. S3B**). Based on these results, we further characterized those 49 most dominant epitopes using predicted binding for additional HLA class II alleles, including a panel of the 12 most common HLA-DQ and DP allelic variants, and all HLA class II variants (DR, DQ, and DP) expressed in the cohort.

Overall, the 49 most dominant epitopes showed significantly higher binding promiscuity (number of alleles bound at the 1,000 nM or better threshold) (Paul et al., 2019; Southwood et al., 1998) for the panel of common HLA class II than a control group of 49 non-epitopes derived from the same proteins (Average number of HLA predicted to be bind ± SD epitopes = 10.8 ± 6.5; non-epitopes = 5.7 ± 6; p=0.0001 by Mann-Whitney; **Fig. S3C-D**). The same conclusion was reached when the full set of HLA alleles present in the cohort were considered using the same criteria (Average ± SD epitopes = 24.3 ± 15.2; non-epitopes = 13.2 ± 14.1; p= 0.0003 by Mann-Whitney; **Fig. S3E-F**).

Heat maps of the 49 epitopes and non-epitopes considering the panel of common HLA DR, DP, and DQ are shown in **Fig. 3.** These results confirm that broad HLA binding capacity is a key feature of dominant epitopes. It further indicates that, because of their broad binding capacity, these epitopes are likely to be recognized in different geographical settings and different ethnicities.

**Fig. 3.**
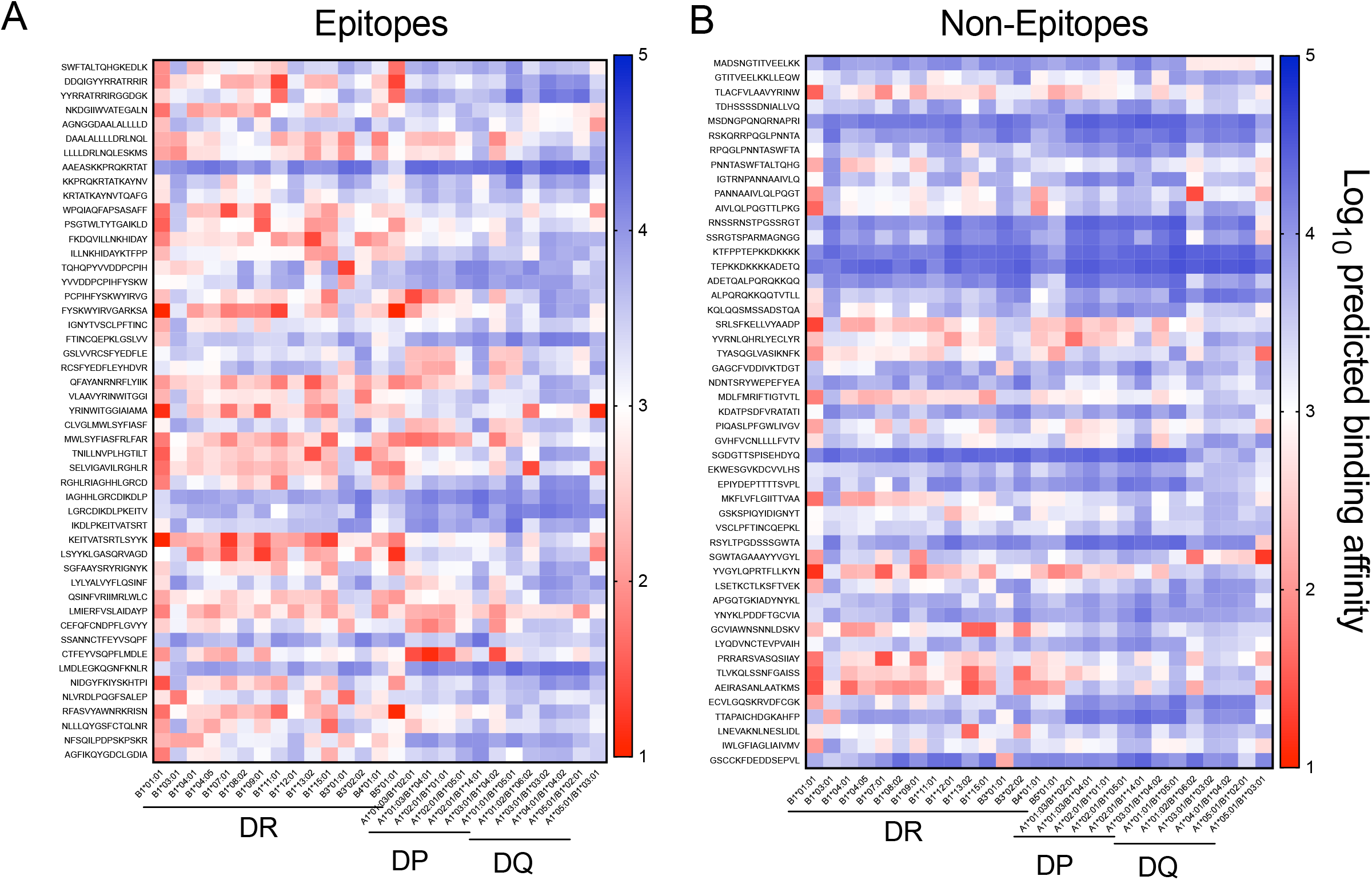
Heat maps of HLA predicted binding patterns in the 27 most frequent HLA class II alleles worldwide (Greenbaum et al., 2011). Predicted binding patterns for the top 49 most immunodominant SARS-CoV-2 CD4^+^ T cell epitopes (**A**) are compared with a set of matched non-epitopes (**B**). Predicted IC_50_ were calculated using NetMHCIIpan embedded in Tepitool (Dhanda et al., 2019; Karosiene et al., 2013; Paul et al., 2016) and converted to Log_10_ scale. Lower values indicate stronger predicted binding affinity, and are highlighted at the red end of the spectrum. Predicted values with an IC_50_ <1000nM (Log_10_ scale <3) are considered positive binders (Paul et al., 2019; Southwood et al., 1998).

### Similarity of SARS-CoV-2 CD4+ T cell epitopes to CCC sequences

Several studies have reported significant preexisting immune memory to SARS-CoV-2 peptides in unexposed donors (Braun et al., 2020; Grifoni et al., 2020; Le Bert et al., 2020; Mateus et al., 2020). This reactivity was shown to be associated, at least in some instances, with memory T cells specific for human common cold coronaviruses (CCC) cross-reactively recognizing SARS-CoV-2 sequences (Braun et al., 2020; Mateus et al., 2020). In particular, it was shown that the SARS-CoV-2 epitopes recognized in unexposed donors had significantly higher homology to CCC than SARS-CoV-2 sequences not recognized in unexposed donors. Here, using the exact same methodology (Mateus et al., 2020), we performed the converse analysis, namely an analysis of the homology between the CD4^+^ T cell epitopes experimentally identified in COVID-19 donors (**Fig. S4**) and sequences of peptides derived from the four widely circulating human CCC (NL63, OC43, HKU1, 229E). No significant differences were observed based on percent sequence identity between epitopes recognized from the COVID-19 cohorts and non-epitope controls in structural proteins S, M, and N (**Fig. S4A**), and non-structural proteins (**Fig. S4B**) or accessory proteins encoded by ORF3a and ORF8 (**Fig. S4C**).

Indeed, in our previous studies (Grifoni et al., 2020; Mateus et al., 2020), we noted that the pattern of antigen recognition in exposed and unexposed donors was significantly different. Here, having defined the actual epitopes recognized in COVID-19, we compared them to the epitopes previously identified in unexposed donors. The present study re-identified 50% of the epitopes in our COVID-19 cohort, but in addition identified 227 novel CD4^+^ T cell epitopes specific for SARS-CoV-2 infection (**Fig. S4G**). Thus, more than 80% (227/280) of the epitopes identified herein are novel and were not previously seen in the unexposed cohort. These results are consistent with the notion that while a cross-reactive repertoire is present in unexposed donors, SARS-CoV-2 infection elicits a vast repertoire of novel T cell specificities.

### Summary of CD8+ T cell epitope identification results

Following the approach described above, a total of 523 SARS-CoV-2 CD8^+^ T cell epitopes were identified (**Table S5**). These epitopes are associated with 26 different HLA restrictions, based on predicted HLA binding capacity matched to the HLA alleles of the responding donor. For eight HLAs, only 1-2 donors expressing the matching HLA could be tested. Predicted binders for the remaining 18 HLAs were tested in a median of 5 donors (range 3 to 9). The 8 most immunodominant proteins were screened in an average of 19 donors (range 4 to 35) (**Fig. 4B**). Of the 523 CD8^+^ T cell epitopes identified, 61 were recognized in 2 or 3 different donor-allele combinations, meaning that there were 454 unique peptides recognized. Of these, 101 (22%) were recognized by 2 or more donors, accounting for 49% of the total response. We found that each donor recognized an average of 2.7 antigens and responded to an average of 1.6 CD8^+^ T cell epitopes per antigen per HLA allele (data not shown) (**Fig. 1F**). Considering 4 HLA A and B alleles in each donor, we expect at least 17 epitopes per donor for class I (2.7 × 1.6 × 4 = 17.3).

**Fig. 4.**
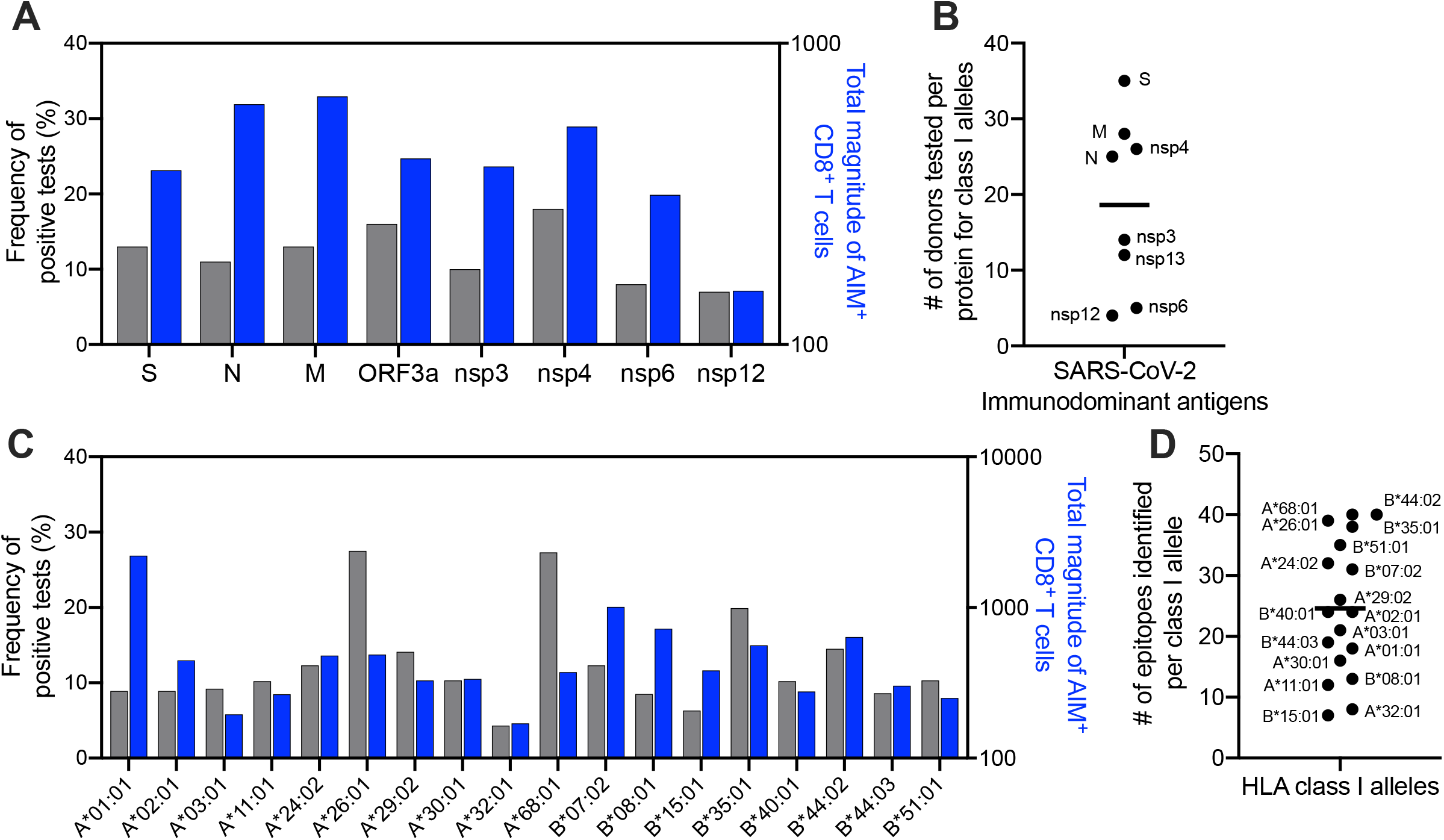
Distribution of allele-specific CD8^+^ responses for the 18 class I alleles that were tested in 3 or more donors as function of protein composition (**A**) or the HLA class I alleles tested (**C**). Blue bars represent the total magnitude of AIM^+^ CD8^+^ T cells divided by the number of positive donors. Gray bars represent the frequency of positive tests. The number of donors tested with each of the 8 dominant proteins for CD8^+^ is shown in panel (**B**). The total number of epitopes identified for each class I allele is shown in panel (**D**).

**Figure 4** shows the frequency of positive epitopes (identified epitopes/peptides screened), and the average magnitude of epitope responses (total magnitude of response normalized by the number of positive donors), as a function of protein (**Fig. 4A**) or HLA class I allele (**Fig. 4C)** analyzed. Each HLA was associated with an average of 25 epitopes (range 7 to 40, median 24) (**Fig. 4D**). Interestingly, as also previously detected in other systems (Goulder et al., 1997; Weiskopf et al., 2013), there was a wide variation as a function of HLA allele. Some alleles, such as A*03:01 and A*32:01, were associated with responses that were both infrequent and weak; in other cases (e.g. A*01:01), responses were infrequent, but when observed were of high magnitude. Finally, and conversely, other alleles were associated with relatively frequent but low magnitude responses (e.g. A*68:01). This effect was previously linked to differences in the size of peptide repertoires associated with different HLA motifs (Paul et al., 2013).

In terms of antigen specificity of CD8^+^ T cell responses, relatively similar epitope-specific response frequencies were observed for the various antigens, with the exception of nsp12, which was associated with responses of low frequency and magnitude (**Fig. 4A**). These results should be interpreted with the caveat in mind that the donors screened were pre-selected on the basis of association with positive responses to that particular antigen; thus, this data does not directly address protein immunodominance, which is instead addressed in **Table 2**. These data instead point to the relative frequency and magnitude of responses at the level of individual epitopes associated with a given antigen, which were found to be overall similar.

To address the potential relationship between CD8^+^ T cell epitope recognition and CCC homology, as performed above in the case of CD4^+^ T cell epitopes, we analyzed the homology of the CD8^+^ T cell epitopes to CCC (NL63, OC43, HKU1, 229E), as compared to the homolog to the same CCC viruses detected in the case of peptides that tested negative in all donors tested, regardless of the HLA-restriction (**Fig. S4**). Similar to what was observed in the context of CD4^+^ T cell responses, the CD8^+^ T cell epitopes recognized in convalescent COVID-19 donors were not associated with higher sequence identity to CCC as compared to non-epitopes, when structural (**Fig. S4D**), accessory (**Fig. S4E**) or non-structural proteins (**Fig. S4F**) were considered.

### Distribution of CD4^+^ and CD8^+^ T cell epitopes within dominant SARS-CoV-2 antigens

We next analyzed the distribution of CD4^+^ and CD8^+^ T cell epitopes within the dominant SARS-CoV-2 S, N, and M antigens (**Fig. 5**). For each antigen, we show the frequency (red line) and magnitude (black line) of CD4^+^ T cell responses along the antigen sequence, considering regions with response frequency above 20% as immunodominant. Based on the results presented above, we also plotted HLA class II binding promiscuity (defined as the number of HLA allelic variants expressed in the donor cohort predicted to be bound by a given peptide), and the degree of homology of each 15-mer peptide for aligned CCC antigen sequences. The bottom panel represents the distribution of CD8^+^ T cell epitopes (black) and non-epitopes (red) along the antigen sequence.

**Fig. 5.**
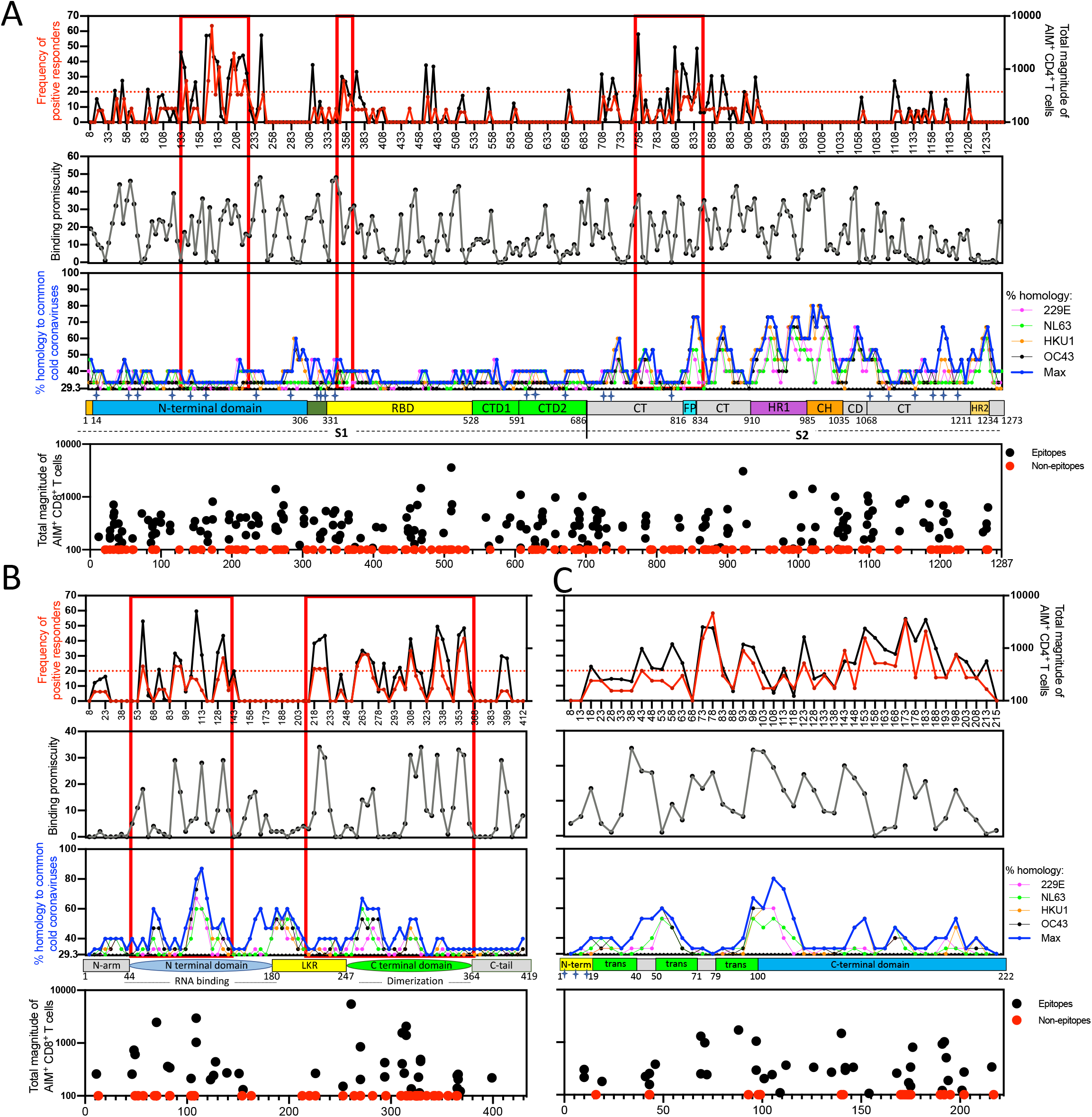
Immunodominant regions for CD4^+^ T cell reactivity for S (**A**), N (**B**) and M (**C**) proteins as a function of the frequency of positive response (red) and total magnitude (black) in the topmost panel. The dotted red line indicates the cutoff of 20% frequency of positivity used to define the immunodominant regions boxed in red and also shown in red in **Fig. 6**. The x-axis labels in this topmost panel indicate the middle position of the peptide. Binding promiscuity was calculated based on NetMHCIIpan predicted IC_50_ for the alleles present in the cohort of donors tested and is shown in grey on the upper middle panel. The lower middle panel shows the % homology of SARS-CoV-2 to the four most frequent CCC (229E in pink, NL63 in green, HKU1 in orange, and OC43 in black) and the max value (blue). The linear structure of each protein is drawn below the graph of homology (Cai et al., 2020; Zeng et al., 2020; UniProtKB - P59596 (VME1_SARS)). The magnitude of CD8^+^ responses to class I predicted epitopes is shown in the bottom panel, where black dots represent epitopes and red dots represent non-epitopes, each centered on the middle position of the peptide.

Responses to S peptides with a frequency of 20% or higher were focused on discrete regions of the protein involving the N-terminal domain (NTD), the C-terminal (CT) 686-816 region, and the neighboring fusion protein (FP) region; only a few responses were focused on the RBD. These immunodominant regions are boxed in red in **Fig. 5A**. We expected HLA-binding capacity to be associated with T cell immunodominant regions, and indeed found a significant positive correlation with the frequency of responses (R = 0.2231, p = 0.0003 by Spearman correlation, **Fig. S5A**). No significant correlation (R = −0.03144, p = 0.6187 by Spearman correlation, **Fig. S5B**) was found with sequence homology to CCC (calculated as maximum sequence homology to the four main CCC species). As indicated in the 3D rendering of the S crystal structure (PDB ID: 6XR8), these immunodominant regions were mostly located in the surface-exposed portions of the S monomer, and were not particularly influenced by the glycosylation pattern (shown in **Fig. 5A** as stars in the linear structure description, and based on experimental identification by Cai and co-authors (Cai et al., 2020). The glycosylation patterns are also shown in the 3D-rendering of the corresponding crystal structure, based on curation done by the authors of the same manuscript, and shown as grey dots (**Fig. 6A**). We further explored the correlation between CD4^+^T cell immunodominance and location of proteolytic cleavage sites, utilizing the MHCII-NP algorithm (Paul et al., 2018). The results did not reveal any significant correlation between the predicted cleavage sites and immunodominant regions (Spearman correlation has R= −0.08426 and p= 0.1816, **Fig. S5C**). This is consistent with previous results that indicated that predicted cleavage sites do not significantly improve epitope predictions (Paul et al., 2018). Finally, CD8^+^ T cell reactivity did not reveal any particular immunodominant region in S, with epitopes and non-epitopes roughly equally distributed along the sequence (**Fig. 5A**).

**Fig. 6.**
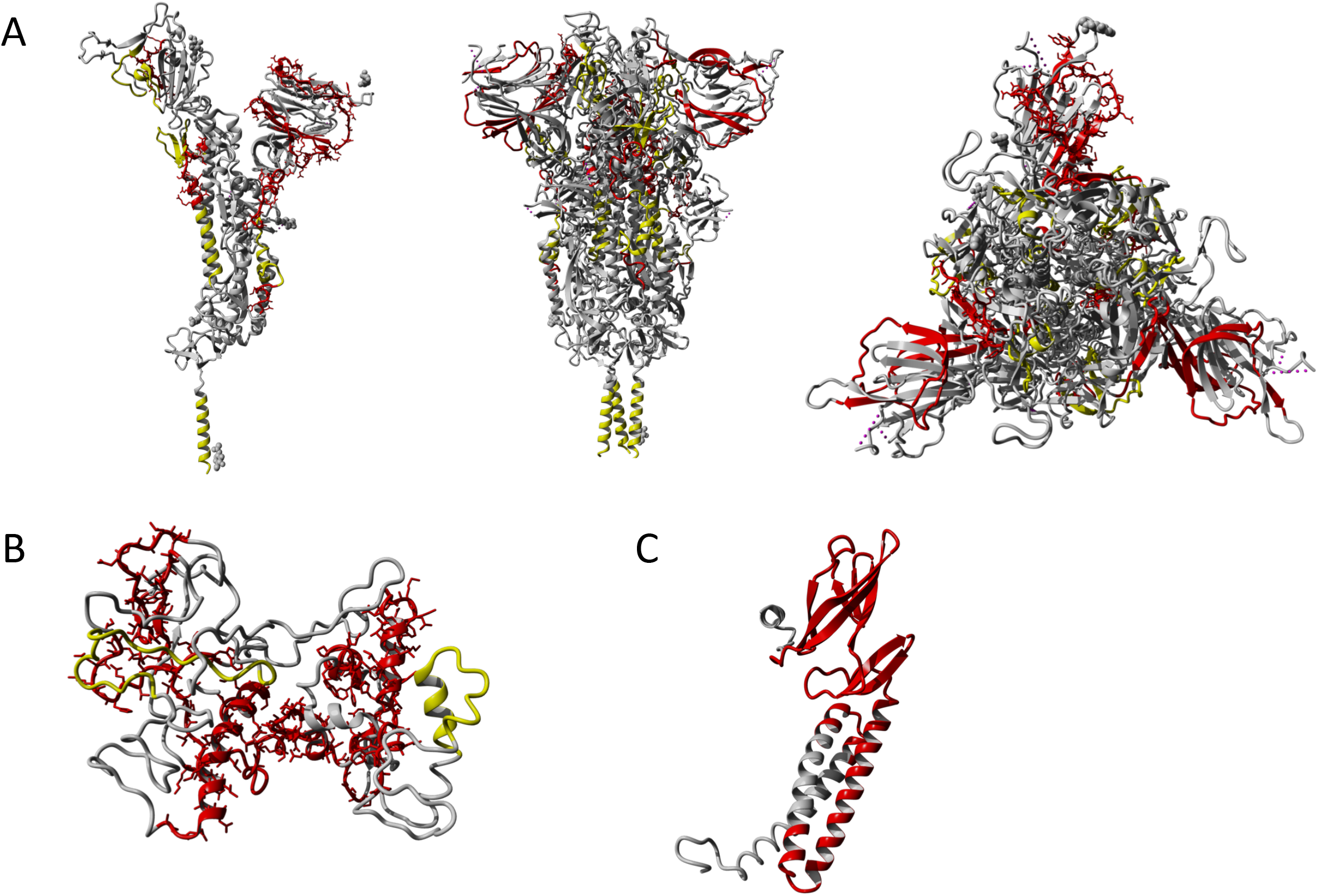
3D-rendering of S (**A**), N (**B**), and M (**C**) proteins. The drawings show in grey the 3D-structures, in red the CD4^+^ T cell immunodominant regions for each protein with frequency of positive responses >20% (also shown in red in **Fig. 5**), and in yellow the B cell immunodominant regions for each protein based on the work of Shrock et al., 2020. Glycosylation sites for S are shown as grey dots and are based on information embedded in the original crystal structure shown to map the immunodominant regions (PDB ID: 6XR8) (**A**) The S protein is shown as monomer on the left and trimer in the middle and on the right (side and top views). (**B**) N protein 3D-rendering was based on a model generated using Phyre2 (Kelley LA et al., 2015). Additional details about the N model are available in the Materials and Methods section. The M protein is shown as a monomer according to a model previously described by Heo et al., 2020 (**C**). All the 3D-rendering have been performed using the free version of YASARA (Land et al., 2018).

In the same way, we compared responses observed within the N and M proteins as a function of structural protein composition, HLA promiscuity and CCC homology (**Fig. 5B-C** and **Fig. 6B-C**). For the N protein (**Fig. 5B**), the majority of the response was focused on the NTD and CTD regions, with lower contributions from the linker region (all outlined in red boxes); segments in the middle and towards the ends of the protein were devoid of any reactivity. The correlation between immunodominance and HLA binding promiscuity was even stronger than observed for S (R = 0.4725, p <0.0001; **Fig. S5D**). Similar to what was observed for the S protein, no significant correlation between the frequency of positive responses was observed with CCC similarity (R = 0.1660, p = 0.1362; **Fig. S5E**) or predicted cleavage sites (R = −0.009245, p = 0.9343; **Fig. S5F**). The immunodominance of N-specific CD8^+^ T cell responses mirrors the one observed for the CD4^+^ T cell counterpart, highlighting that in general the N-terminal and C-terminal domains are the major immunodominant regions of N recognized by both T cell types.

CD4^+^ T cell immunogenic regions were distributed across the entire span of the M protein **(Fig. 5C)**, including the transmembrane region (**Fig. 6C**). No significant correlation was observed when investigating HLA binding promiscuity (R = 0.2374, p = 0.1253; **Fig. S5G**), CCC similarity (R = 0.07648, p = 0.6259; **Fig. S5H**), or predicted cleavage sites (R = 0.08421, p = 0.5913; **Fig. S5I**). The lack of correlation between M epitopes and HLA binding is consistent with the interpretation that M is a prominent antigen because it is highly expressed, not because it contains high quality epitopes. No particular immunodominance patterns were observed for the M protein with respect to CD8^+^ epitopes.

Finally, when we investigated the location of immunodominant T cell regions relative to the main sites identified for antibody reactivity (Shrock et al., 2020), the CD4^+^ T cell immunodominant regions identified in S and N showed minimal overlap with immunodominant linear regions targeted by antibody responses (Shrock et al., 2020) (**Fig. 6**). The CD4^+^ T cell epitope recognition patterns of ORF3a, ORF8, nsp3, nsp4, nsp12, and nsp13 are shown in **Fig S6.**The ORF8 protein was similar to M in that epitopes throughout both of these small proteins were recognized. ORF3a had clear regions of response clustered in the middle and at the C-terminus. Nsp3, which was the 4^th^ most immunodominant antigen, was associated with a rather striking immunodominant region centered around residue 1643. Other non-structural proteins were less immunodominant overall, but had discreet regions targeted by CD4^+^ T cell responses (i.e. residue 5253 for nsp12).

### Reactivity of megapools based on the experimentally identified epitopes

The experiments described above identified a total of 280 CD4^+^ and 454 CD8^+^ T cell epitopes. These epitopes were arranged into two epitope megapools (MPs), CD4-E and CD8-E, respectively (where the E denotes “experimentally defined”). These MPs were tested in a new cohort of 31 COVID-19 convalescent donors (none of these donors were utilized in the epitope identification experiments) and 25 unexposed controls (**Table 3**). MP reactivity was assessed for all donors using AIM and IFNγ FluoroSpot assays.

**Table 3.**
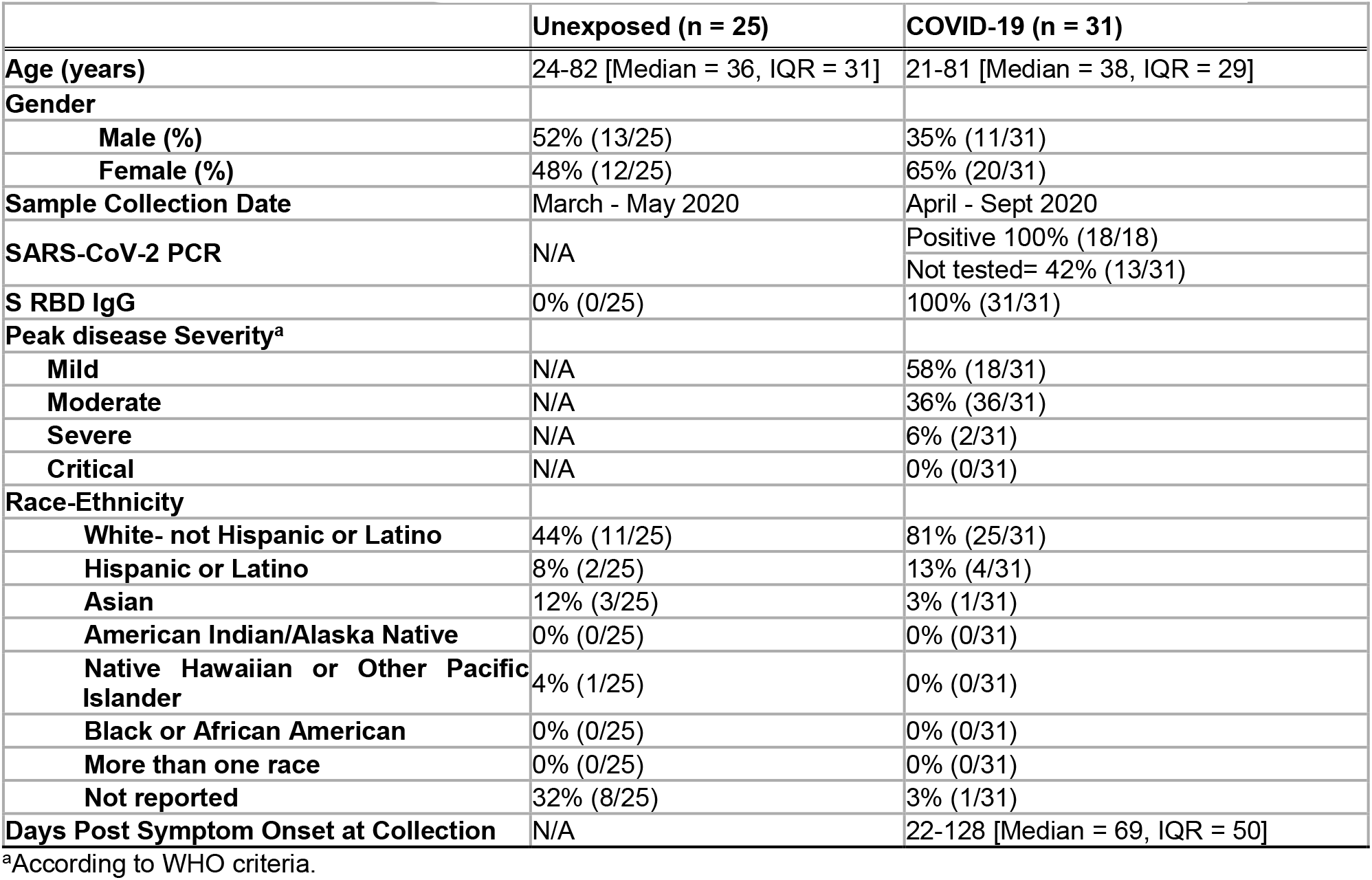
Characteristics of donor cohorts utilized to validate megapools in **Fig. 7**.

To put the results in context, we also tested peptides contained in the CD4-R and CD4-S, and CD8-A and CD8-B MPs previously utilized to measure SARS-CoV-2 CD4^+^ and CD8^+^ T cell responses, respectively (Grifoni et al., 2020; Mateus et al., 2020; Rydyznski Moderbacher et al., 2020; Weiskopf et al., 2020). These MPs are based on either overlapping peptides spanning the entire S sequence (CD4-S) or predicted peptides (all other proteins). While these pools contain a larger total number of peptides (474 for CD4-R+ CD4-S, and 628 for the CD8-A+ CD8-B) than the corresponding experimentally defined sets, we expected that the experimentally defined peptide sets would be able to recapitulate the reactivity observed with the previously utilized MPs. As a further context, we also tested the T cell Epitope Compositions (EC) class I and EC class II pools of experimentally defined CD8^+^ and CD4^+^ epitopes described by Nelde et al. (Nelde et al., 2020), encompassing 29 and 20 epitopes each, which prior to this study represented the most comprehensive set of experimentally defined epitopes.

As might be expected, the results showed that the AIM assay was more sensitive than the FluoroSpot assay (**Fig. 7**). On the other hand, as a tradeoff for the lower signal, the FluoroSpot assay showed higher specificity in the responses detected, with fewer unexposed individuals showing any reactivity compared to the AIM assay. For CD4^+^ T cell responses as detected in the AIM assay (**Fig. 7A**), the CD4-E MP recapitulated the reactivity observed with the MPs of larger numbers of predicted peptides (CD4-R+S), and showed significantly higher reactivity (p= 4.30×10^−6^ by Mann-Whitney) as compared to the EC class II pool. A similar picture was observed when the FluoroSpot assay was utilized (**Fig. 7B**), with a significantly higher reactivity of the CD4-E MP compared to the CD4-R+S (p= 0.0208 by Mann-Whitney), and to the EC class II pool (p= 1.39×10^−7^ by Mann-Whitney). In both AIM and FluoroSpot assays, the CD4-E MP showed the highest capacity to discriminate between COVID-19 convalescent and unexposed donors (p= 3.19×10^−10^ and p= 1.56×10^−9^, respectively by Mann-Whitney).

**Fig. 7.**
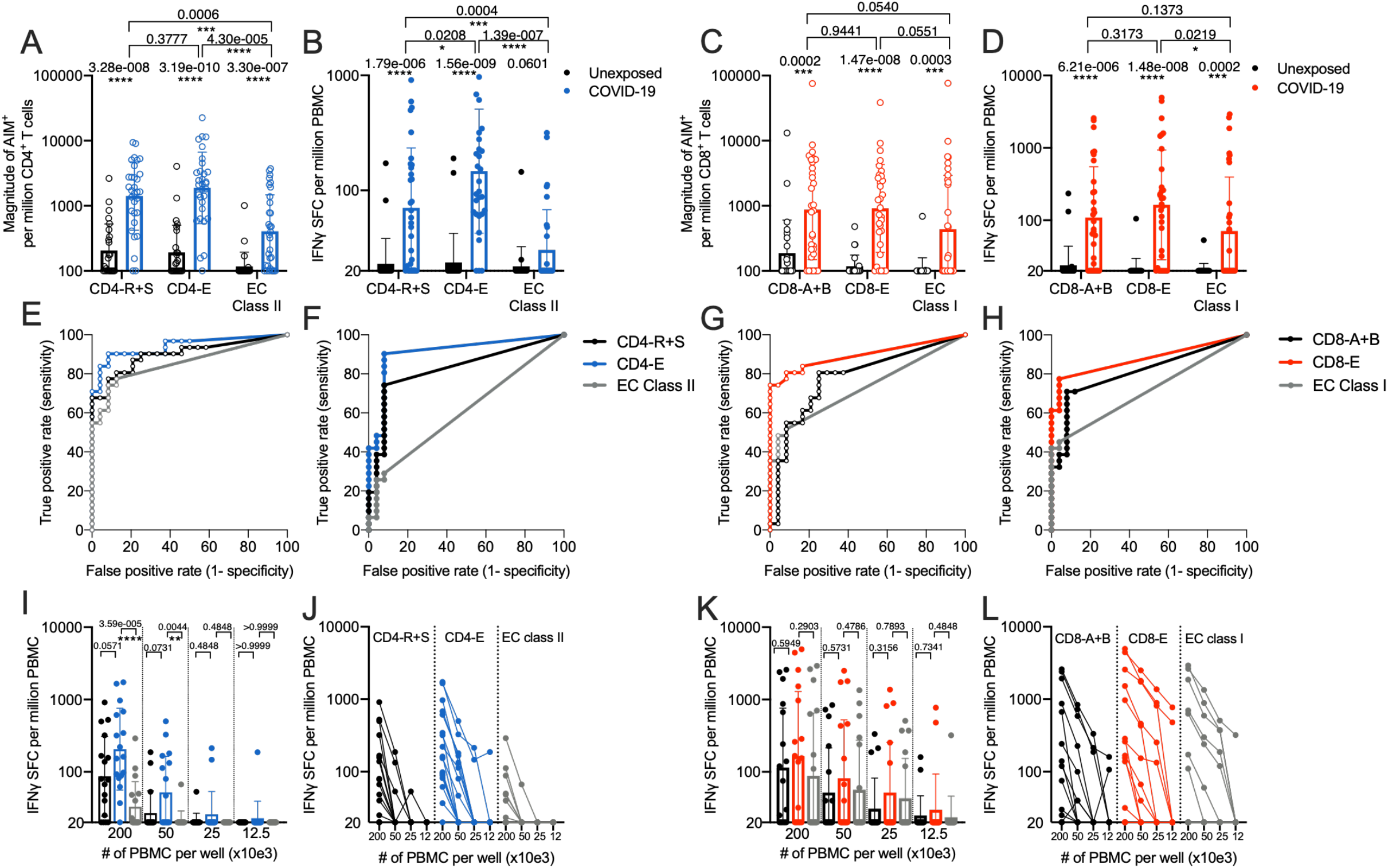
T cell responses to SARS-CoV-2 megapools as measured in AIM (empty circles) and FluoroSpot (filled in circles) assays. Twenty-five unexposed and 31 convalescent COVID-19 donors were tested in the AIM assays (**A**and **C**), and all donors were also tested in the FluoroSpot assays (**B** and **D**). CD4^+^ T cell responses to CD4-R+S (previously described), CD4-E (280 class II epitopes identified in this study), and EC Class II (Nelde et al 2020) megapools were measured via AIM (**A**) and FluoroSpot (**B**). CD8^+^ T cell responses to CD8-A+B (previously described), CD8-E (454 class I epitopes identified in this study), and EC Class I (Nelde et al 2020) megapools were measured via AIM (**C**) and FluoroSpot (**D**). Bars represent geometric mean ± geometric SD, and p-values were calculated by Mann-Whitney. Panels **E-H** show ROC analysis for CD4^+^ and CD8^+^ T cell response data in FluoroSpot (**F-H**) and AIM (**E-G**) assays. In each panel, curves are shown for the 3 peptide pools tested. For a given pool, T cell responses were used to classify individuals into ‘predicted exposed’ or ‘predicted unexposed’, at varying thresholds starting with the highest observed response to the lowest. We then compared these data with the actual SARS-CoV-2 exposure status of the individuals and calculated the rate of true positive (predicted exposed / total exposed) and the rate of false positives (predicted exposed / total non-exposed). Additionally, we further tested 17 of these COVID-19 convalescent donors in FluoroSpot with a titration of 200, 50, 25, and 12.5×10^3^ cells per well with the indicated CD4-MPs (**I-J**) and CD8-MPs (**K-L**).

A similar picture was noted in the case of CD8^+^ T cell reactivity (**Fig. 7C-D**), where the CD8-E MP recapitulated the reactivity observed with the MPs of larger numbers of predicted peptides (CD8-A+B), with a strong trend (p= 0.0551 by Mann-Whitney) towards more reactivity than the EC class II pool. In the case of the FluoroSpot assay, we noted equivalent reactivity for the CD8-E and CD8-A+B MPs, and significantly higher reactivity (p= 0.0219 by Mann-Whitney) than the EC class II pool (**Fig. 7D**). In both assays, the CD8-E MP showed highest capacity to discriminate between COVID-19 convalescent and unexposed subjects (p= 1.47×10^−8^ and p= 1.48×10^−8^, respectively by Mann-Whitney). To test how well the different T cell responses measured separate individuals that have been exposed to SARS-Cov-2 versus those that do not, we performed ROC analysis (**Fig. 7E-H**) which allow to directly compare the classification success based on true-and false-positive rates. The CD4-E and CD8-E response data were associated with the best performance.

Considering that a potential practical limitation in the characterization of SARS-CoV-2 responses is the number of cells available for study, in selected COVID-19 donors we titrated the number of PBMC/well to determine if a response could be measured with lower cell numbers. As expected, as the cell input was decreased, the magnitude of responses decreased correspondingly. While marginal responses were seen with 25,000 cells/well and below, a sizeable response was still detectable with 50,000 cell/well, with 8 out of 17 donors responding for the CD4-E MP (as compared to 16 out of 17 in the case of 200,000 cell level). Similarly, in the case of the CD8-E MP, with 8 out of 17 donors responding (as compared to 11 out of 17 in the case of 200,000 cell level). The frequency and magnitude of responses of CD4-E were higher compared to the EC class II (p= 3.59×10^−5^ and p= 0.0044 by Mann-Whitney) (**Fig. 7I-J**). The CD8-E MP was also associated with a higher magnitude of response than the EC class I pool (**Fig. 7K-L**). In conclusion, these results underline the biological relevance of the more comprehensive CD4-E and CD8-E MPs.

## DISCUSSION

This study presents a comprehensive analysis of the patterns of epitope recognition associated with SARS-CoV-2 infection in humans. The analysis was performed using a cohort of approximately 100 different convalescent donors spanning a range of peak COVID-19 disease severity representative of the observed distribution in the San Diego area. SARS-CoV-2 was probed using 1,925 different overlapping peptides spanning the entire viral proteome, ensuring an unbiased coverage of the different HLA class II alleles expressed in the donor cohort. For HLA class I we used an alternative approach, selecting 5,600 predicted binders for 28 prominent HLA class I alleles, representing 61% of the HLA A and B allelic variants in the worldwide population, and affording an overall 98.8% HLA class I coverage at the phenotypic level.

The biological relevance of the epitope characterization studies summarized here is underlined by the use of the *ex vivo* AIM assay that does not require *in vitro* stimulation, which potentially skews the results by eliciting responses from naïve cells. The AIM assay is also more agnostic for different types of CD4^+^ T cells, as it measures all activated cells, regardless of T cell subset or any particular pattern of cytokine secretion.

We are not aware of any study that describes the repertoire of CD4^+^ and CD8^+^ T cell epitopes recognized in SARS-CoV-2 infection with a comparable level of granularity or breadth. While several previous reports have described SARS-CoV-2 epitopes, and accordingly represent very useful advances, these studies either utilized *in vitro* expansion (Nelde et al., 2020), were limited in the number of proteins analyzed (Le Bert et al., 2020), characterized responses in fewer than 10 HLA types (Ferretti et al., 2020; Nelde et al., 2020; Peng et al., 2020), or focused on TCR repertoire after *in vitro* expansion of small number of cells (Snyder et al., 2020). Comparing our results with those obtained in those previous studies, we note that of the 20 HLA class II peptides identified by Nelde and co-authors(Nelde et al., 2020), 14 were contained within proteins we mapped here in detail, and we independently re-identified 12 (86%) of them (identical or largely overlapping sequences). Of 137 class I peptides reported thus far (Ferretti et al., 2020; Nelde et al., 2020; Peng et al., 2020), 98 were contained within the viral proteins we mapped in detail, and we independently re-identified 68 (69%) of them (identical or largely overlapping sequences).

Importantly, because SARS-CoV-2 antigen-specific T cell responses were evaluated in a systematic and unbiased fashion, quantitative estimates of the size of the repertoire of T cell epitope specificities recognized in each donor can be derived. Determining the breadth of responses is of relevance, since previous studies (Ferretti et al., 2020; Snyder et al., 2020) have suggested narrow SARS-CoV-2-specific T cell repertoires in COVID-19 patients; notably, a limited repertoire could favor viral mutation, a particular concern with this RNA virus. Based on our results, we expect that each donor would be able to recognize about 19 CD4^+^ T cell epitopes, on average. Likewise, for CD8^+^ T cells, we expect at least 17 epitopes per donor to be recognized. Overall, T cell responses in SARS-CoV-2 are estimated to recognize even more epitopes per donor than seen in the context of other RNA viruses, such as dengue (Grifoni et al., 2017; Weiskopf et al., 2015), where 11.6 and 7 CD4^+^ and CD8^+^ T cell epitopes, respectively, were recognized on average. This analysis should allay concerns over the potential for SARS-CoV-2 to escape T cell recognition by mutation of a few key viral epitopes.

We defined the patterns of immunodominance across the various antigens encoded in the SARS-CoV-2 genome recognized in COVID-19 donors. Consistent with earlier reports from our group (Grifoni et al., 2020) and others (Peng et al., 2020), we see clear patterns of immunodominance, with a limited number of antigens accounting for about 80% of the total response. In general, the same antigens are dominant for both CD4^+^ and CD8^+^ responses, with some differences in relative ranking, such as in the case of nsp3, which is relatively more dominant for CD8^+^ than CD4^+^ T cell responses. Immunodominance at the protein level correlated with predicted expression levels, as previously noted for CD4^+^ T cell responses, although we note that the accessory proteins and nsps also account for a significant fraction of the response despite their predicted lower abundance in infected cells.

Because of their role in instructing both antibody and CD8^+^ T cell responses, we correlated CD4^+^ T cell activity on a per donor and per antigen level with antibody and CD8^+^ T cell adaptive responses. This enabled establishing which antigens have functional relevance in terms of eliciting CD4^+^ T cell responses correlated with antibody and CD8^+^ T cell responses. At the level of antibody responses, S and M were correlated with RBD antibody titers, highlighting their capacity to support antibody responses, presumably by a deterministic linkage (viral antigen bridge) and cognate interactions (Sette et al., 2008). Surprisingly, N-specific CD4^+^ T cell responses did not correlate with S RBD antibody titers, suggesting unexpected complexity of the N-specific CD4^+^ T cell response. By contrast with these selective effects, CD4^+^ T cell activity against any of the antigens correlated with the total CD8^+^ T cell activity, suggesting that the role of CD4^+^ T cell responses driven by the different proteins is determinant in its helper function for either RBD-specific antibody production or CD8^+^ T cell responses. This was particularly true in both contexts when looking specifically at the S and M proteins, which are also the strongest and most frequently recognized antigens for both CD4^+^ and CD8^+^ T cells.

After examining relative immunodominance at the level of the different SARS-CoV-2 antigens, we probed for variables that may influence which specific peptides are recognized within a given antigen/ORF. Previously, we have shown that SARS-CoV-2 sequences recognized in unexposed individuals were associated with a higher degree of similarity to sequences encoded in the genome of various CCC. Here, repeating the same analysis with the SARS-CoV-2 epitopes recognized in COVID-19 donors, we found no significant correlation. We further show that while a large fraction of the epitopes previously identified in unexposed donors are re-identified in COVID-19 donors, about 80% of the epitopes are novel (not previously seen in unexposed), suggesting that the SARS-CoV-2-specific T cell repertoire of COVID-19 cases is overlapping, but substantially different from, the SARS-CoV-2-cross-reactive memory T cell repertoire of unexposed donors. This is consistent with our previous observation of a different pattern of reactivity (Mateus et al., 2020), and consistent with reports from other groups (Le Bert et al., 2020; Nelde et al., 2020).

HLA binding capacity was a major determinant of immunogenicity for CD4^+^ T cells (the influence of HLA binding was not evaluated for CD8^+^ T cell, since the tested epitope candidates were picked based of their predicted HLA binding capacity). As found in several previous large-scale pathogen-derived epitope identification studies, immunodominant epitopes were also found to be promiscuous HLA class II binders (Lindestam Arlehamn et al., 2016; Oseroff et al., 2010). Binding to multiple HLA allelic variants is an important mechanism to amplify the potential immunogenicity of peptide epitopes and specific regions within an antigen. It is possible that the dominance of particular regions might further correlate with processing. However, at this juncture, HLA class II processing algorithms do not effectively predict epitope recognition (Barra et al., 2018; Cassotta et al., 2020; Paul et al., 2018).

Further analysis projected the CD4^+^ T cell dominant regions on known or predicted SARS-CoV-2 protein structures. This established that the dominant epitope regions are different for B and T cells. This is of relevance for vaccine development, as inclusion of antigen sub-regions selected on the basis of dominance for antibody reactivity might result in an immunogen devoid of sufficient CD4^+^ T cell activity. In this context, it is important to note that the RBD region had very few CD4^+^ T cell epitopes recognized in COVID-19 donors, but inclusion of regions neighboring the RBD N-and C-termini would be expected to provide sufficient CD4^+^ T cell help.

In contrast to the clear demarcation of dominant regions for antibody and CD4^+^ T cell responses, CD8^+^ T cell epitopes were uniformly dispersed throughout the various antigens, consistent with previous in-depth analyses revealing little positional effect in CD8^+^ T cell epitope distribution (Kim et al., 2013). In the case of CD8^+^ T cell responses, our data highlights HLA-allele specific differences in the frequency and magnitude of responses. This effect was noted before in the case of dengue virus (Weiskopf et al., 2013) and related to potential HLA-linked protective versus susceptibility effects. The current study is not powered to test these potential effects, leaving it to future studies to examine this possibility. Regardless, our study provides a roadmap for inclusion of specific regions or discrete epitopes, to allow for CD8^+^ T cell epitope representation across a variety of different HLAs.

Finally, the functional relevance of our study was highlighted by the generation of novel and improved epitope MPs for measuring T cell responses to SARS-CoV-2; these newer experimentally defined pools are associated with increased activity and lower complexity when compared to our previous MPs based on overlapping and predicted peptides. We plan to make these epitope pools available to the scientific community at large, and expect that they will facilitate further investigation of the role of T cell immunity in SARS-CoV-2 infection and COVID-19.

In conclusion, we identify several hundred different HLA class I and class II restricted SARS-CoV-2-derived epitopes. We anticipate that these results will be of significant value in terms of basic investigation of SARS-CoV-2 immune responses, in the development of multimeric staining reagents, and in the development of T cell-based diagnostics. In addition, the results shed light on the mechanisms of immunodominance of SARS-CoV-2, which have implications for understanding host-virus interactions, as well as for vaccine design.

### Limitations and future directions

To maximize cell usage, our analysis was focused on the most dominantly recognized proteins. Screening for less commonly recognized proteins would require a larger cohort to enable identification of a sufficient number of donors responding to each protein. However, such expanded studies would be expected to yield additional epitopes.

The limited number of donors studied also did not allow investigation of responses directed against relatively rare HLA alleles, and HLA restrictions were not experimentally verified. The predictions utilized for HLA class I included the top 200 candidates for each allele. Utilizing more generous prediction thresholds is likely to allow for identification of additional epitopes. The limited number of donors also did not allow for the evaluation of potential differences in terms of ethnic background, disease severity, age, and gender. Future investigations will include validation of the epitope pools as potential diagnostic tools, establish a robust, user-friendly T cell assay, and investigate differences in T cell reactivity as a function of ethnicity, disease severity, age, and gender.

## ACKNOWLEDGEMENTS

This study has been funded by the NIH NIAID (award AI42742 to S.C., A.S., contract Nr. 75N9301900065 to A.S. and D.W., NIH grant U01 CA260541-01 to DW, K08 award AI135078 to J.D., and AI036214 to D.S.). Additional support has been provided by UCSD T32s (AI007036 and AI007384 to S.A.R and S.I.R) and the Jonathan and Mary Tu Foundation (D.S.). A.T. was supported by a PhD student fellowship through the Clinical and Experimental Immunology Course at the University of Genoa, Italy. We thank Gina Levi and the LJI clinical core for assistance in sample coordination and blood processing, and Erica Ollmann Saphire, Michael Norris, and Sara Landeras-Bueno for useful discussions and input on 3-D modeling.

## AUTHOR CONTRIBUTIONS

Conceptualization: A.T., A.G., S.C. and A.S.; Data curation and bioinformatic analysis, J.A.G. and B.P.; Formal analysis: A.T., J.S., C.K., A.G., D.W., J.M.D, J.M. E.W.; Funding acquisition: S.C., A.S., D.W., S.I.R., S.A.R., and J.M.D.; Investigation: A.T., E.W., C.K., N.M., J.M.D, J.M., J.S., E.M., P.R., D.W. A.S and A.G.; Project administration: A.F. Resources: S.I.R., S.A.R., A.C., S.M., E.P., D.M.S., S.C., and A.S.; Supervision: B.P., J.S., A.d.S., S.C., D.W., R. d. A., A.S., and A.G.; Writing: A.T., S.C., A.S., and A.G.

## DECLARATION OF INTEREST

A.S. is a consultant for Gritstone, Flow Pharma, Merck, Epitogenesis, Gilead and Avalia. S.C. is a consultant for Avalia. All other authors declare no conflict of interest. LJI has filed for patent protection for various aspects of vaccine design and identification of specific epitopes.

## STAR METHODS

### RESOURCE AVAILABILITY

#### Lead Contact

Further information and requests for resources and reagents should be directed to the lead contact, Dr. Alessandro Sette.

#### Materials Availability

Epitope pools used in this study will be made available to the scientific community upon request, and following execution of a material transfer agreement, by contacting Dr. Alessandro Sette.

#### Data and Code Availability

The published article includes all data generated or analyzed during this study, and summarized in the accompanying tables, figures and supplemental materials.

### EXPERIMENTAL MODEL AND SUBJECT DETAILS

#### Human Subjects

##### Convalescent COVID-19 Donors utilized for epitope identification

Blood donations from the 99 convalescent donors included in this study’s cohort were collected through either the UC San Diego Health Clinic under IRB approved protocols (200236X), or under IRB approval (VD-214) at the La Jolla Institute. Donations obtained through the CROs Sanguine, BioIVT and Stem Express were collected under the same IRB approval (VD-214) at the La Jolla Institute. Details of this cohort can be found in **Table 1**. All donors were over the age of 18 years and no exclusions were made due to disease severity, race, ethnicity, or gender. All donors were able to provide informed consent, or had a legal guardian or representative able to do so. Study exclusion criteria included lack of willingness or ability to provide informed consent, or lack of an appropriate legal guardian to provide informed consent.

Disease severity was defined as mild, moderate, severe or critical as previously described (Grifoni 2020). In brief, this classification of disease severity is based on a modified version of the WHO interim guidance, “Clinical management of severe acute respiratory infection when COVID-19 is suspected” (WHO Reference Number: WHO/2019-nCoV/clinical/2020.4). At the time of enrollment in the study, 80% of donors had been confirmed positive by swab test viral PCR during the acute phase of infection. Plasma samples from all donors were later tested by IgG ELISA for SARS-CoV-2 S protein RBD to verify previous infection (**Table 1 and Fig. 2A**).

##### Healthy Unexposed donors utilized for CD4-E and CD8-E megapool validation

Samples from healthy adult donors were obtained from the San Diego Blood Bank (SDBB). According to the criteria set up by the SDBB if a subject was eligible to donate blood, they were considered eligible for our study. All the donors were tested for SARS-CoV-2 RBD IgG serology and were found negative and therefore considered unexposed. An overview of the characteristics of these donors is provided in **Table 3**.

##### Convalescent COVID-19 donors utilized for CD4-E and CD8-E megapool validation

The 31 convalescent donors tested in the megapool AIM and FluoroSpot assays (**Fig. 7**) were collected from the same clinics using the same protocols as described above for the donors utilized for epitope identification. Similarly, no donors enrolled were under the age of 18 and none were excluded due to disease severity, race, ethnicity, or gender. All donors, or legal guardians, gave informed consent. Specific characteristics of these donors can be found in **Table 3**, including the summary of ELISA testing for SARS-CoV-2 S protein RBD.

### METHOD DETAILS

#### Peptide Pools

##### Preparation of 15-mers and subsequent megapools and mesopools

To identify SARS-CoV-2-specific T cell epitopes, 15-mer peptides overlapping by 10 amino acids and spanning entire SARS-CoV-2 proteins were synthesized. All peptides were synthesized as crude material (A&A, San Diego, CA) and individually resuspended in dimethyl sulfoxide (DMSO) at a concentration of 10 mg/mL. Aliquots of these peptides were pooled by antigen of provenance into megapools (MP) (as described in **Table 2**) and sequentially lyophilized as previously reported (Carrasco Pro et al., 2015). Another portion of the 15-mer peptides were pooled into smaller mesopools of ten peptides each. All pools were resuspended at 1 mg/mL in DMSO.

##### Class I peptide preparation

Class I predicted peptides were designed using the protein sequences derived from the SARS-CoV-2 reference strain (GenBank: MN908947). Predictions were performed as previously reported using NetMHC pan EL 4.0 algorithm (Jurtz et al., 2017) for 28 HLA A and B alleles that were selected based on frequency in our cohort and also representative of the worldwide population (**Fig. S1A-B**). The top 200 predicted peptides were selected for each allele. In total 5,600 class I peptides were synthesized and resuspended in DMSO at 10 mg/mL.

#### PBMC isolation and HLA typing

Whole blood was collected from all donors in either Acid Citrate Dextrose (ACD) tubes or heparin coated blood bags. Whole blood was then centrifuged at room temperature for 15 minutes at 1850 rpm to separate the cellular fraction and plasma. The plasma was then carefully removed from the cell pellet and stored at −20C. Peripheral blood mononuclear cells (PBMC) were isolated by density-gradient sedimentation using Ficoll-Paque (Lymphoprep, Nycomed Pharma) as previously described (Weiskopf et al., 2013). Isolated PBMC were cryopreserved in cell recovery media containing 10% DMSO (Gibco), supplemented with 90% heat-inactivated fetal bovine serum, depending on the processing laboratory, (FBS; Hyclone Laboratories, Logan UT) and stored in liquid nitrogen until used in the assays. Each sample was HLA typed by Murdoch University in Western Australia, an ASHI-Accredited laboratory (Voic 2020, Madden 1995, Gorse 2010). Typing was performed for the class I HLA A and B loci and class II DRBI, DQB1, and DPB1 loci.

#### SARS-CoV-2 RBD ELISA

The SARS-CoV-2 RBD ELISA has been described in detail elsewhere (Grifoni 2020, Amanat 2020). All convalescent COVID-19 donors had their serology determined by ELISA. Briefly, 96-well half-area plates (ThermoFisher 3690) were coated with 1 ug/mL SARS-CoV-2 Spike (S) Receptor Binding Domain (RBD) and incubated at 4°C overnight. On the following day plates were blocked at room temperature for 2 hours with 3% milk in phosphate buffered saline (PBS) containing 0.05% Tween-20. Then, heat-inactivated plasma was added to the plates for another 90-minute incubation at room temperature followed by incubation with conjugated secondary antibody, detection, and subsequent data analysis by reading the plates on Spectramax Plate Reader at 450 nm using SoftMax Pro. Limit of detection (LOD) was defined as 1:3. Limit of sensitivity (LOS) for SARS-CoV-2 infected individuals was established based on uninfected subjects, using plasma from normal healthy donors not exposed to SARS-CoV-2.

#### Flow Cytometry

##### Activation induced cell marker (AIM) assay

The AIM assay was performed as previously described (Dan et al., 2016; Reiss et al., 2017). Cryopreserved PBMCs were thawed by diluting the cells in 10 mL complete RPMI 1640 with 5% human AB serum (Gemini Bioproducts) in the presence of benzonase [20μl/10ml]. Cells were cultured for 20 to 24 hours in the presence of SARS-CoV-2 specific MPs [1 μg/ml], mesopools [1 μg/ml], 15-mers [10 μg/ml], or class I predicted peptides [10 μg/ml] in 96-wells U bottom plates with 1×10^6^ PBMC per well. As a negative control, an equimolar amount of DMSO was used to stimulate the cells as a negative control in triplicate wells, and phytohemagglutinin (PHA, Roche, 1μg/ml) was included as the positive control. The cells were stained with CD3 AF700 (4:100; Life Technologies Cat# 56-0038-42), CD4 BV605 (4:100; BD Biosciences Cat# 562658), CD8 BV650 (2:100; Biolegend Cat# 301042), and Live/Dead Aqua (1:1000; eBioscience Cat# 65-0866-14). Activation was measured by the following markers: CD137 APC (4:100; Biolegend Cat# 309810), OX40 PE-Cy7 (2:100; Biolegend Cat#350012), and CD69 PE (10:100; BD Biosciences Cat# 555531). All samples were acquired on either a ZE5 cell analyzer (Bio-rad laboratories) or an Aurora flow cytometry system (Cytek), and analyzed with FlowJo software (Tree Star).

#### HLA binding assays

The binding of selected SARS-CoV-2 15-mer epitopes to HLA class II MHC molecules was measured as previously described (Sidney 2013, Voic 2020). In brief, the binding is quantified by each peptide’s capacity to inhibit the binding of a radiolabeled peptide probe to purified MHC in classical competition assays. The probe was incubated with purified MHC, a mixture of protease inhibitors, and different concentrations of unlabeled inhibitor peptide at room temperature or 37°C for 2 days. MHC molecules were subsequently captured on HLA-DR-specific monoclonal antibody (L243) coated Lumitrac 600 plates (Greiner Bio-one, Frickenhausen, Germany) and radioactivity was measured using the TopCount microscintillation counter (Packard Instrument Co., Meriden, CT). Each peptide was tested at 6 concentrations to cover a 100,000-fold dose range, and an unlabeled version of the radiolabeled probe was included in each experiment as a positive control for inhibition. To analyze the results, we calculated the concentration of peptide at which the binding was inhibited by 50% (IC50 nM). For these values to approximate true Kd values, the following conditions were met: 1) the concentration of radiolabelled probe is less than the concentration of MHC, and 2) the measured IC50 is greater than or equal to the concentration of MHC.

#### FluoroSpot

PBMCs derived from 25 unexposed donors were stimulated in triplicate at a single density of 200×10^3^ cells/well (one donor was tested at 50×10^3^ due to limitation in cell numbers). PBMCs from a cohort of 31 convalescent COVID-19 donors were stimulated in triplicates of 200×10^3^ cells/well, with the exception of 5 donors tested at 50-100×10^3^ cells/well due to cell limitations (**Fig. 7B, D, F,** and **H**). Seventeen of these convalescent donors were further titrated at 200, 50, 25, and 12.5×10^3^ cells/well (**Fig. 7I-L**). The cells were stimulated with the different MPs analyzed (1µg/mL), PHA (10µg/mL), and DMSO (0.1%) in 96-well plates previously coated with anti-cytokine antibodies for IFNγ, (mAbs 1-D1K; Mabtech, Stockholm, Sweden) at a concentration of 10µg/mL. After 20 hours of incubation at 37°C, 5% CO_2_, cells were discarded and FluoroSpot plates were washed and further incubated for 2 hours with cytokine antibodies (mAbs 7-B6-1-BAM; Mabtech, Stockholm, Sweden). Subsequently, plates were washed again with PBS/0.05% Tween20 and incubated for 1 hour with fluorophore-conjugated antibodies (Anti-BAM-490). Computer-assisted image analysis was performed by counting fluorescent spots using an AID iSPOT FluoroSpot reader (AIS-diagnostika, Germany). Each megapool was considered positive compared to the background based on the following criteria: 20 or more spot forming cells (SFC) per 10^6^ PBMC after background subtraction for each cytokine analyzed, a stimulation index (S.I.) greater than 2, and statistically different from the background (p<0.05) in either a Poisson or T test.

### BIOINFORMATIC AND STATISTICAL ANALYSIS

FlowJo 10 and GraphPad Prism 8.4 were used to perform data and statistical analyses, unless otherwise stated. Statistical details of the experiments are provided in the respective figure legends. Data plotted in linear scale are expressed as mean + standard deviation (SD). Data plotted in logarithmic scales are expressed as median + 95% confidence interval (CI) or geometric mean + geometric SD. Statistical analyses were performed using Spearman correlation and Mann-Whitney or Kolmogorov-Smirnov tests for unpaired comparisons. Details pertaining to significance are also noted in the respective figure legends.

#### AIM assay analysis

In analyzing data from the AIM assays, the counts of AIM^+^ CD4^+^ and CD8^+^ T cells were normalized based on the counts of CD4^+^ and CD8^+^ T cells in each well to be equivalent to 1×10^6^ total CD8^+^ or CD4^+^ T cells. The background was removed from the data by subtracting the single or the average of the counts of AIM^+^ cells plated as single or triplicate wells stimulated with DMSO. We included the triplicate wells stimulated with DMSO in the mesopools and epitope identification steps to take into account the variability of the weaker signals observed in those two respect to the original MP reactivity (da Silva Antunes et al., 2020). The Stimulation Index (SI) was calculated by dividing the count of AIM^+^ cells after SARS-CoV-2 stimulation with the ones in the negative control. A positive response had an SI greater than 2 and a minimum of 100 AIM^+^ cells after background subtraction. The gates for AIM^+^ cells were drawn relative to the negative and positive controls for each donor. A representative example of the gating strategy is depicted in **Fig. S2B**.

#### HLA class I nested epitopes

For some alleles and proteins, multiple nested class I predicted peptides were tested in the AIM assay. In cases where a specific donor responded to multiple nested epitopes corresponding to the same allele and protein, the epitope with the highest magnitude of response was classified as the optimal epitope. If multiple nested epitopes had the same response (within a range of 50 AIM^+^ cells), the epitope with the shortest length was selected. Nested epitopes corresponding to different donors or different alleles were conserved as separate epitopes.

#### CCC homology analysis

SARS-CoV-2-derived 15-mer peptides were analyzed for their identity with the common cold coronaviruses (CCC) 229E, NL63, HKU1, and OC43, as previously described (Mateus et al., 2020). In brief, every SARS-CoV-2 15-mer peptide tested for immunogenicity was compared against every position in the corresponding protein sequences of common coronaviruses obtained from GenBank. The region that best matched the respective SARS-CoV-2 peptide was used to calculate percent sequence identity for each of the four CCC viruses individually, as well as the maximum across all four (**Fig. S5**). The same methodology was also used to calculate sequence identity for SARS-CoV-2 class I peptides (**Fig. S5**). Using the same set of common coronavirus reference sequences, an alternative analysis was performed by mapping each SARS-CoV-2 peptide with the S, M and N protein sequences corresponding to the four common coronavirus using Immunobrowser tool (Dhanda et al., 2018). The values resulted from this specific analysis are plotted in **Fig. 5**.

#### T cell epitope restriction predictions

Putative HLA class II restrictions for individual 15-mer CD4^+^ T cell epitopes were inferred using the IEDB’s TepiTool resource (Paul 2016). All CD4^+^ T cell prediction analyses were performed applying the NetMHCIIpan algorithm (Karosiene et al., 2013). Prediction analyses were performed to either infer HLA restriction based on the HLA typing of the cohort (**Table S1** and **Table S3**) or to assess potential binding promiscuity of experimentally defined epitopes, considering the 27 most frequent class II alleles in the worldwide population (Greenbaum et al., 2011). In both types of prediction analyses, a 20th percentile threshold was applied (**Table S3**), as previously described (Mateus et al., 2020).

#### Assigning regions within the linear structure

Simple diagrams were created to describe the linear structures of S, N, and M proteins (**Fig. 4**). The different regions of the S protein were defined based on the works of Cai et al. 2020. The structure of the N protein was divided into 3 main regions, the N-and C-terminal domains, and the linker region in between (Zeng et al., 2020). For the M protein, the regions of the structure were extracted from UniProt (UniProtKB - P59596 (VME1_SARS).

#### 3D-rendering and model design

Three different approaches have been used to map T and B cell immunodominant regions on the 3D-structures for SARS-CoV-2 S, M and N proteins. The S protein model was based on the crystal structure described in Cai et al. 2020 (PDB ID: 6XR8) and using the glycosylation sites annotated in the submitted PDB. The M protein model has been previously described by Heo et al., 2020. The model for the N protein was run on four different homology prediction servers (SWISS-MODEL, RaptorX, iTasser and Phyre2). In order to have a complete N sequence, Phyre2 server was subsequently selected using the intensive mode (Kelley and Sternberg, 2009). The resulting model showed a variable level of confidence with higher percentages (>90%) in the C-Terminal domain (CTD) and N-terminal domain (NTD) regions and low confidence percentages (>10%) in the linker domain. The N model was superimposable with both the crystal structures for the CTD (PDB ID: 6WZO) and NTD (PDB ID: 6M3M). The current N model has the only purpose of visualization for mapping immunodominant regions. All the mapping analyses have been performed using the free version of YASARA (Land and Humble, 2018).

## SUPPLEMENTARY MATERIALS

### SUPPLEMENTARY FIGURE LEGENDS

**Fig. S1.**
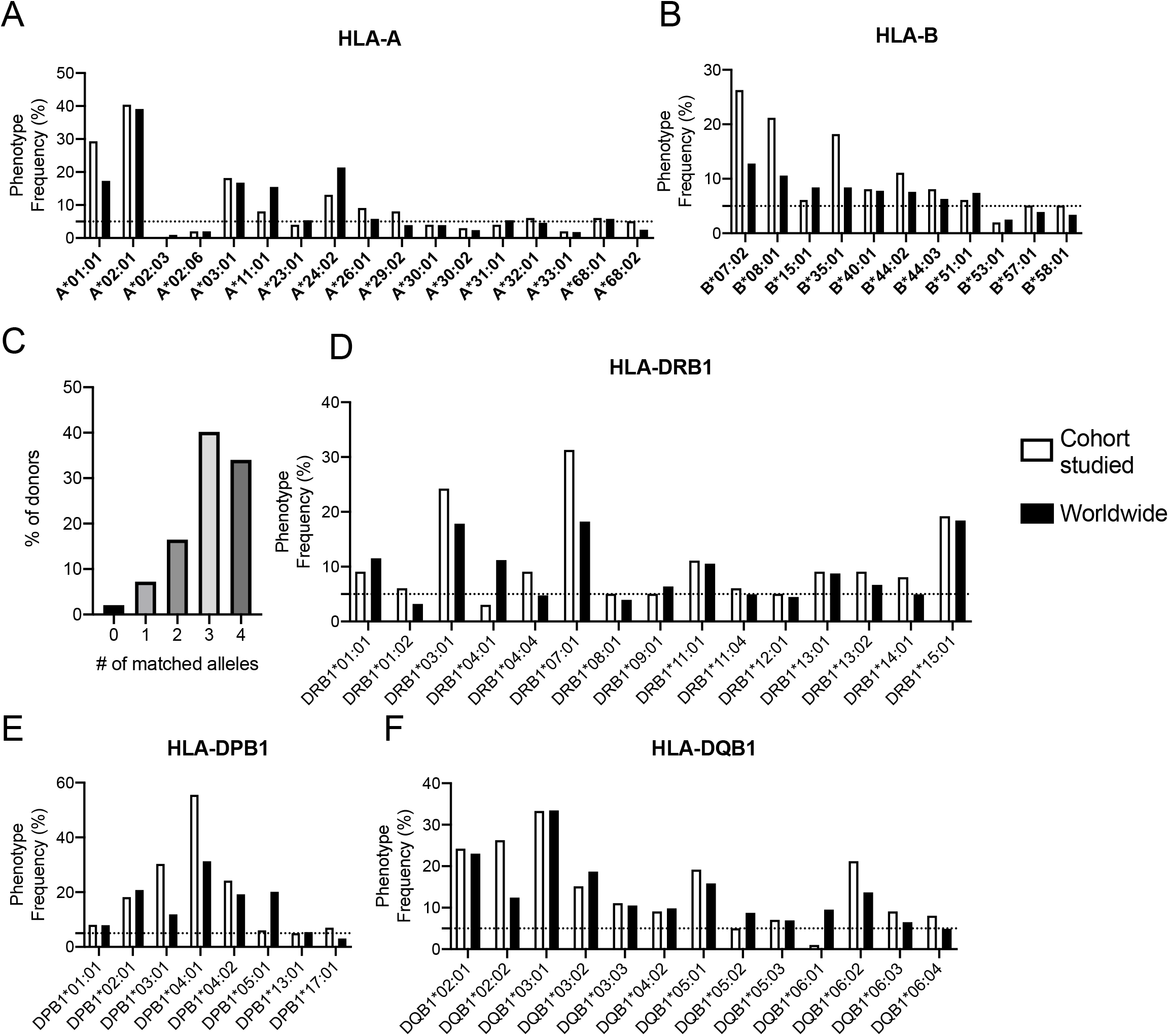
Refers to **Fig. 4** and **Fig. 5**. HLA phenotype frequency in the COVID-19 cohort analyzed compared with the worldwide phenotype frequencies available in the IEDB-AR population coverage tool (Bui et al., 2006; Dhanda et al., 2019). HLA class I frequency for A and B loci for the top 28 HLA class I with frequency >5% in the worldwide population are shown in panels **A** and **B**, respectively. (**C**) Coverage of class I predicted peptides based on the HLA typing of the population. HLA class II frequency for DRB1, DP and DQ loci for the top HLA class II with frequency >5% in the worldwide population or the studied cohort are shown in panels **D**, **E**, and **F** respectively.

**Fig. S2.**
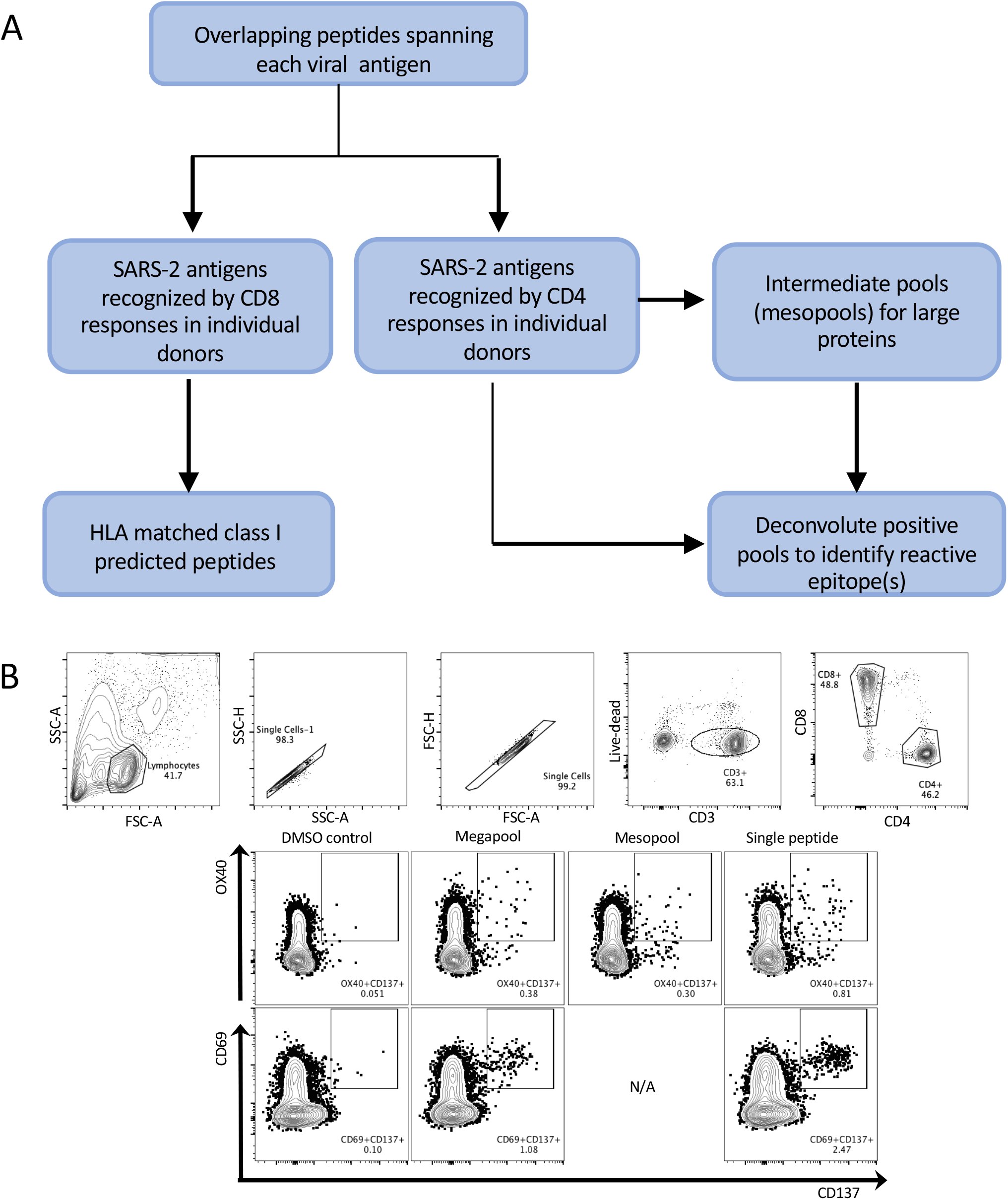
Refers to **Fig. 1**. Summary of experimental strategy. (**A**) Scheme of experimental strategy selected for HLA class I and class II epitope identification. (**B**) Representative graphs depicting the flow cytometry gating strategy for defining antigen-specific CD4^+^ and CD8^+^ T cells by OX40^+^CD137^+^ and CD69^+^CD137^+^ expression, respectively.

**Fig. S3.**
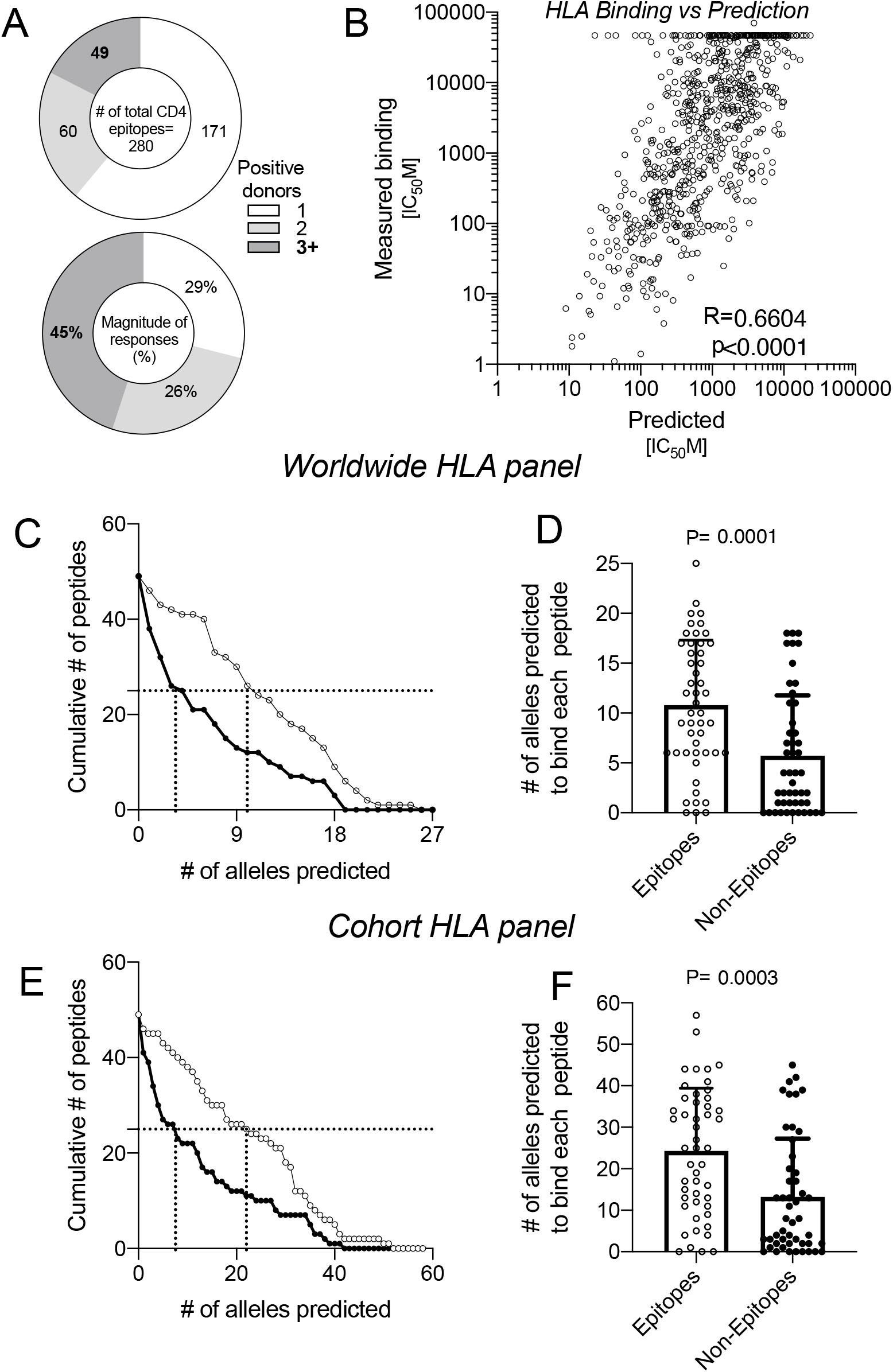
Refers to **Fig. 3**. SARS-CoV-2 immunodominant epitope HLA class II binding capacity and promiscuity. (**A**) CD4 SARS-CoV-2 epitopes as function of the number of responding donors recognized and strength of responses. These data highlight that 49 immunodominant epitopes account for 45% of the total response. A comparison of the HLA class II binding capacity of 49 immunodominant epitopes as determined by binding predictions or as measured experimentally (**B**), suggesting feasibility for using binding predictions to assess HLA-restriction. Predicted HLA class II binding promiscuity is shown for the same 49 epitopes (white circles), and also 49 non-epitopes (black circles), considering the 27 HLA class II alleles most frequent worldwide (**C-D**), or the 58 HLA class II alleles specific to the study cohort (**E-F**).The number of HLA class II alleles predicted to bind epitopes (white circles) and non-epitopes (black circles) are based on a prediction cutoff value of IC_50_≤1000nM. Statistical comparisons were performed using Mann-Whitney.

**Fig. S4.**
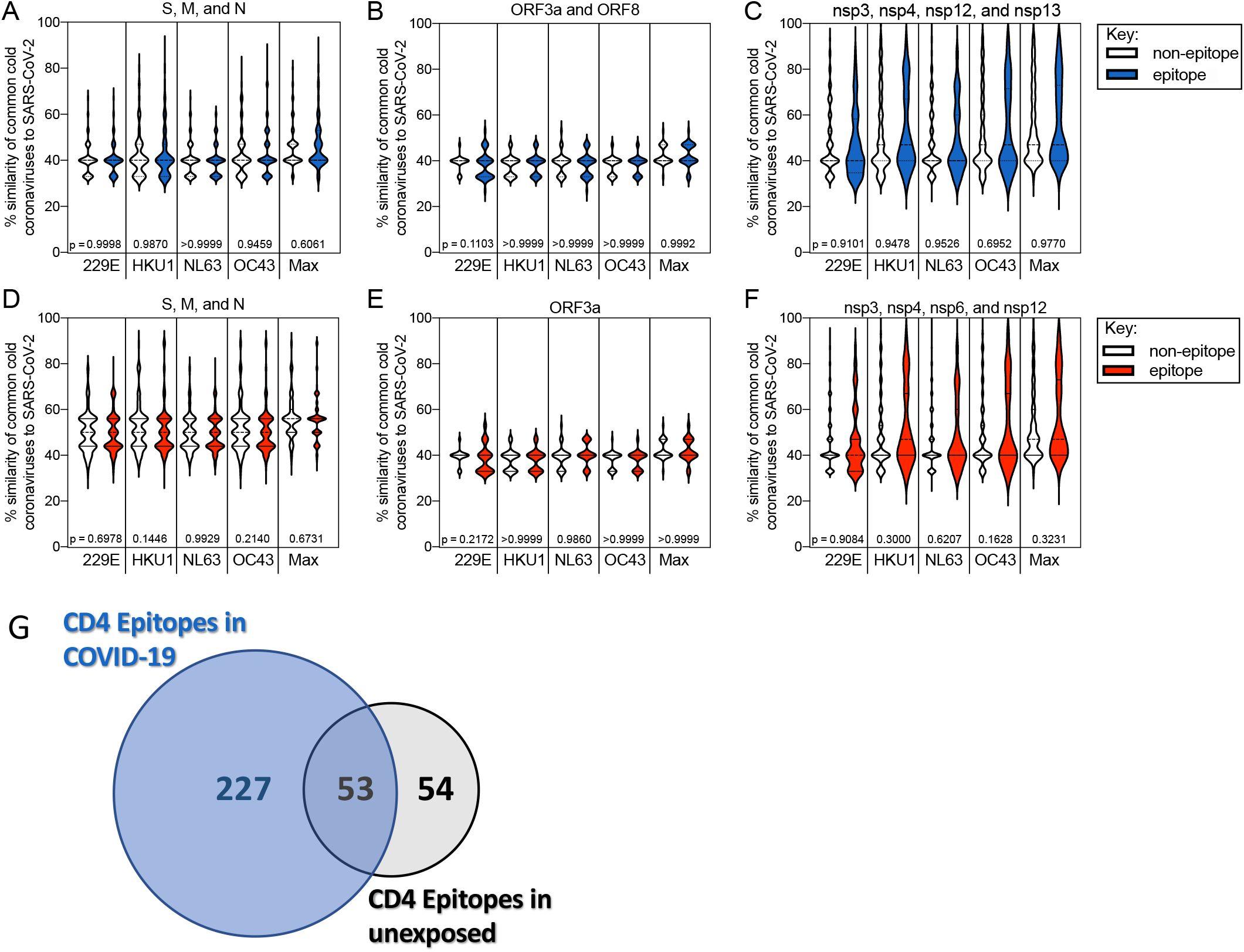
Refers to **Fig. 4** and **5**. Analyses of CD4^+^ and CD8^+ +^ T cell epitopes identified compared to non-epitopes within the same proteins. Comparison of sequenced identity between CD4^+^ T cell epitopes and non-epitopes as a function of sequence identity with the CCC in S, M, and N combined (**A**), ORF8 and ORF3a (**B**), and non-structural proteins (**C**). For CD8^+^ epitopes and non-epitopes, the sequence identities with CCC are shown for S, M, and N (**D**), ORF3a (**E**), and non-structural proteins (**F**). Statistical analyses were performed using the Kolmogorov-Smirnov test, and data are shown as violin plots. (**G**) Overlap of previously identified epitopes in unexposed (Mateus et al., 2020 Science) with the proteins analyzed in this study and the current epitopes identified in COVID-19 donors. The Venn diagram was calculated with the Venn Diagram Plotter (PNNL, OMICS.PNL.gov).

**Fig. S5.**
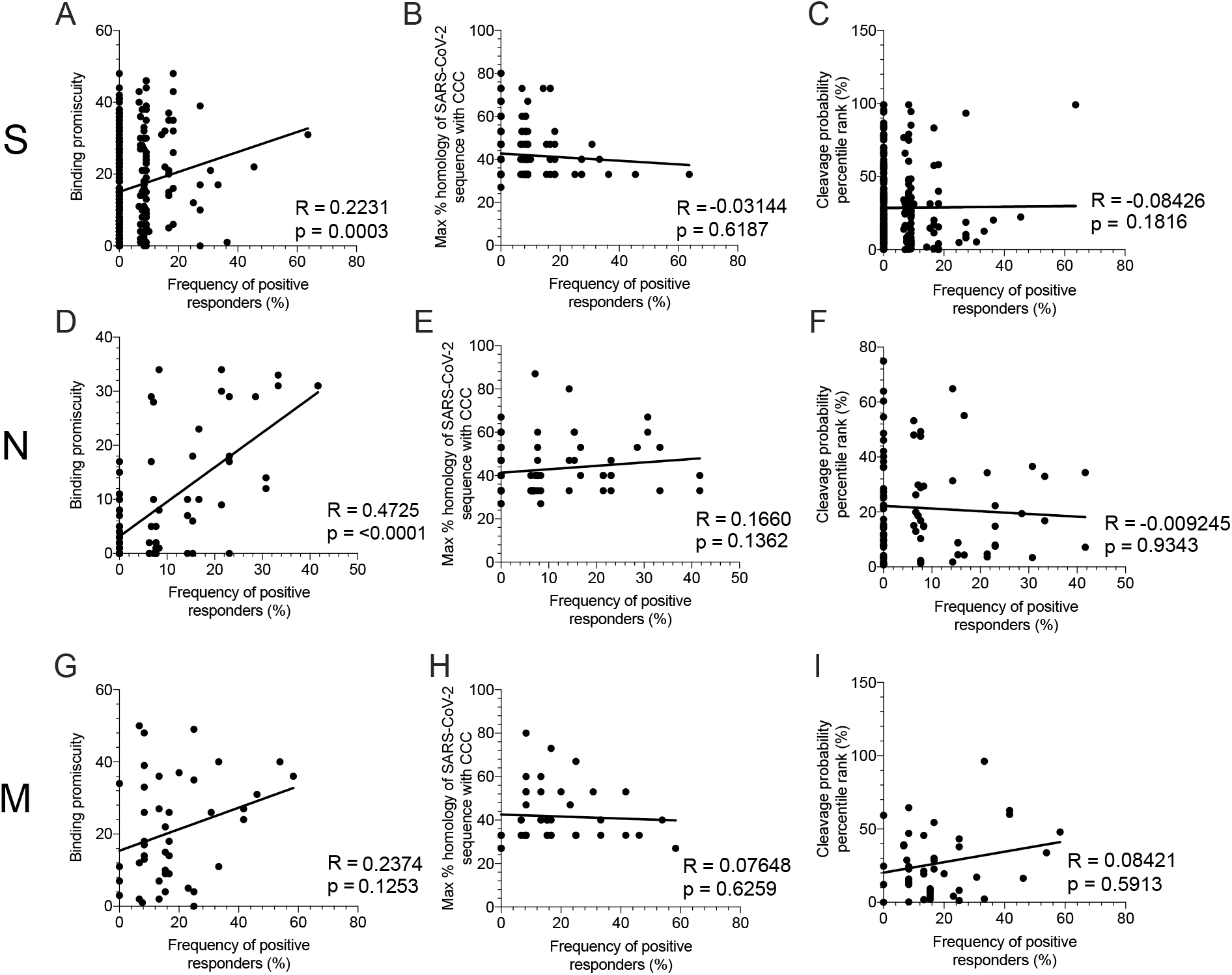
Refers to **Fig. 5**. Correlations of predicted binding promiscuity to the alleles present in the donor cohort tested with the frequency of positive response for S (**A**), N (**D**), and M (**G**) epitopes. Frequency of positive response is also correlated with the maximum % homology of the SARS-CoV-2 sequence to CCC and plotted for S (**B**), M (**E**), and N (**H**). In the final column of panels, the correlation of frequency of positivity and the cleavage probability percentile rank (calculated using the IEDB MHCII-NP tool) are shown for S (**C**), N (**F**), and M (**I**). Statistics were performed using the Spearman correlation and the line on each graph is a simple linear regression.

**Fig S6.**
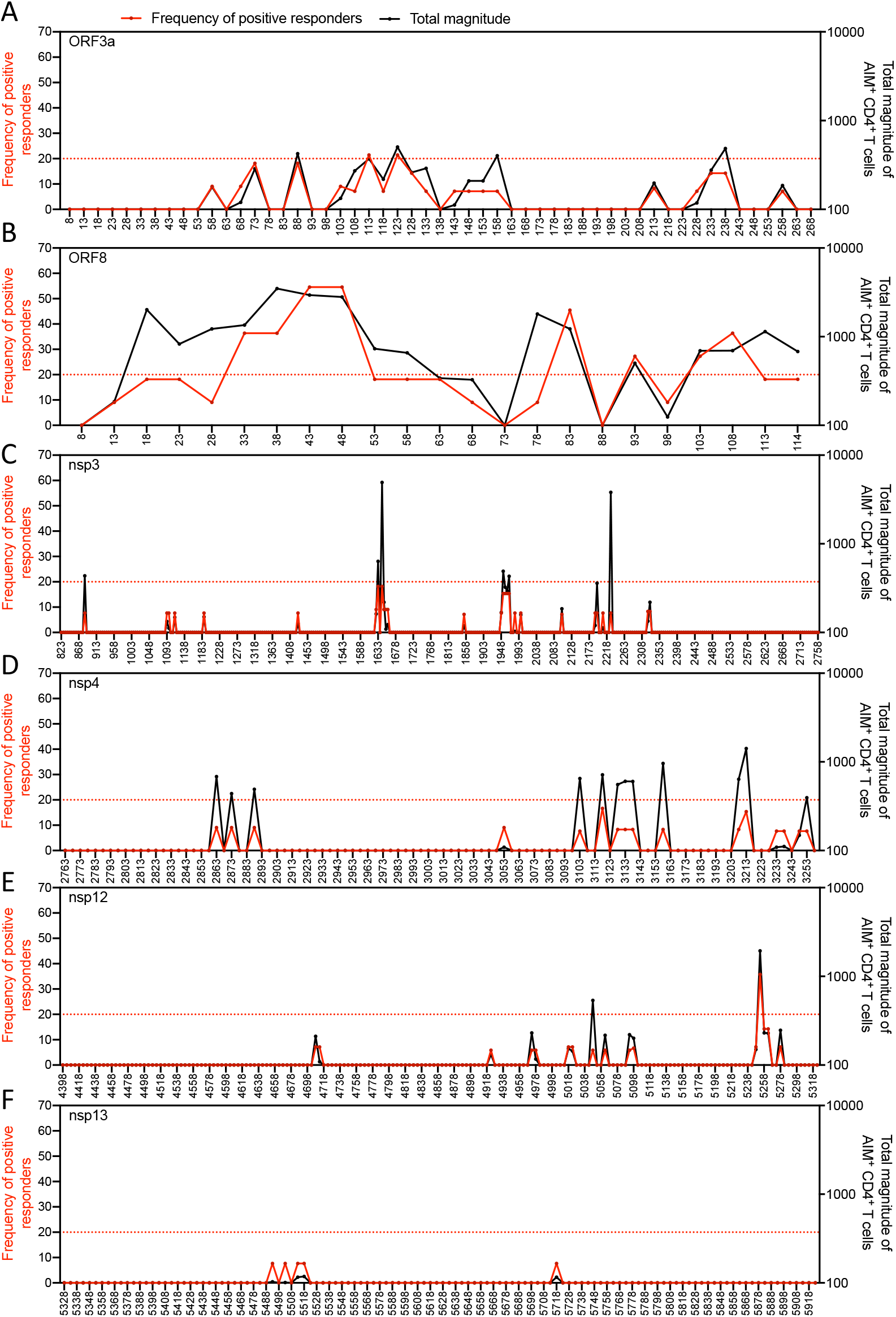
Refers to **Fig. 5**. Immunodominant regions for the other major antigens for CD4^+^ T cells: ORF3a (**A**), ORF8 (**B**), nsp3 (**C**), nsp4 (**D**), nsp12 (**E**), and nsp13 (**F**). The frequency of positive responders is shown in red and the total magnitude of response (sum of AIM^+^ T cells for each peptide) is shown in black. The x-axis is labeled with the middle, or 8^th^, position of each 15-mer peptide. The dotted red line at 20% frequency of positive responders indicates a cutoff for immunodominant epitopes.

### SUPPLEMENTARY TABLE LEGENDS

**Table S1.** HLA typing for class I and class II molecules in the donor cohort. Predicted epitopes were synthesized based on the most frequent 28 HLA class I alleles in the general population. The donors selected for further testing for class I and/or class II epitope identification are indicated.

**Table S2.** Number of predicted epitopes synthesized based on the most frequent 28 HLA class I alleles. 200 epitopes were synthesized for each allele to cover the major SARS-CoV-2 antigens.

**Table S3.** List of CD4^+^ T cell epitopes identified in this study and their predicted HLA restriction(s). A total of 280 15-mer epitopes were identified by AIM assay and encompassed the 9 dominant SARS-CoV-2 antigens for CD4^+^ T cells.

**Table S4.** HLA binding analysis of the 49 class II epitopes identified in 3 or more donors.

**Table S5.** List of CD8^+^ T cell epitopes identified in this study and the HLA restrictions. A total of 523 class I epitopes were identified by AIM assay and encompassed the 8 dominant SARS-CoV-2 antigens for CD8^+^ T cells.

